# Fab Dimerization Crosslinks and Activates the γδ T-Cell Receptor

**DOI:** 10.64898/2026.10.08.757522

**Authors:** Zezhi Li, Yihua Fan, Jing Li, Yuanyuan Zhao, Ligang Chen, Yiqing Chen, Zhiming Wang, Yumei Zheng, Yuxuan Zhang, Jun Wang, Hao Wu, Di Wu, Longfei Wang

**Author notes:** These authors contributed equally: Zezhi Li, Yihua Fan, Jing Li, Yuanyuan Zhao.

## Abstract

The gamma delta T-cell receptor (γδ TCR) mediates γδ T-cell responses that are crucial for immune defense, tissue surveillance, and homeostasis. While recent studies suggest that γδ TCRs engage ligands multivalently, how multivalent engagement activates the γδ TCR remains elusive. Here, we present the structures of the Vγ9Vδ2 (clone G115) TCR in both the inactive and the agonist-crosslinked active states. In the γδ TCR lipoprotein complex, cholesterol and phospholipids form a “barrier” that isolates CD3ζζ from the other subunits and likely prevents autoactivation. As an agonist, the Fab fragment of the UCHT1 antibody dimerizes upon binding the Vγ9Vδ2 TCR, thereby crosslinking and activating the receptor. Cryo-EM analysis of Fab-crosslinked Vγ9Vδ2 TCR identifies three Fab dimer interfaces, one between the two light chains and the other two between light and heavy chains. These Fab-Fab interfaces, together with the Fab-CD3ε interface, mediate the assembly of the Vγ9Vδ2 TCR and UCHT1 Fab into 2:2 tetramers. Upon activation, the Vγ9Vδ2 TCR becomes conformationally plastic and loses enriched lipids, including cholesterol and phosphatidylethanolamine. Our findings reveal a dimerization-driven activation mechanism, inform the design of rigid TCR-crosslinking agonists, and suggest multivalency-induced clustering as a general paradigm for γδ TCR activation.

## Introduction

γδ T-cells play an important role in innate and acquired immunity and are associated with various diseases, such as cancer, infection, and inflammation (1, 2). The γδ T-cell receptor (γδ TCR) consists of a TCRγδ heterodimer which is generated by V(D)J combinations and responsible for antigen recognition. The fully-assembled γδ TCR complex also includes three CD3 dimers (CD3δε, CD3γε, CD3ζζ) that are required for transmembrane signaling (3). In contrast to the short cytoplasmic C-termini of TCRγδ, the CD3 subunits have long cytoplasmic tails containing signaling motifs, such as ITAM and proline-rich sequences (PRS) to signal downstream cellular responses (4-6).

The activation mechanisms of γδ TCR and αβ TCR are distinct, despite their similar subunit compositions. In contrast to the major histocompatibility complex (MHC)-dependent antigen recognition of the αβ TCR, the γδ TCR recognizes a broad repertoire of peptide and non-peptide antigens in an MHC-independent manner, including stress-induced antigens and phosphate antigens (2). Despite extensive efforts, the activation mechanism of the γδ TCR remains unclear. Recent studies have reported the structures of the γδ TCR complex and its antigen-bound assemblies (7-10). However, the molecular mechanism underlying Fab-induced γδ TCR activation remains unclear.

Here, we report cryo-EM structures of the human γδ TCR in both its apo (ligand/agonist-free) and agonist-bound states. These structures reveal striking differences from the αβ TCR, particularly the extensive flexibility of the antigen-binding (extracellular) domains of the TCRγδ subunits. Integrating these data with mass spectrometry (MS), we characterized the lipids associated with the γδ TCR and their dynamics. Notably, we observed the dimerization of an agonist Fab upon binding to the γδ TCR and elucidated the underlying activation mechanism. Our findings shed light on the activation of the γδ TCR and offer a foundation for future therapeutics development.

## Results

### Structure of a γδ TCR Complex

γδ T cells are classified into three major subpopulations based on the TCRδ chain in their T-cell receptor (Vδ1-3) (2). The Vδ2-expressing subset is the main group of γδ T cells in the peripheral blood, accounting for 50-90% of the total γδ T cells and primarily co-expressed with the Vγ9 chain to form Vγ9Vδ2 T cells (11, 12).

To characterize the γδ TCR in vitro, we selected Vγ9Vδ2 TCRs derived from the G115 clonotype (13). For simplicity, these G115-derived Vγ9Vδ2 TCRs are referred to as Vγ9Vδ2 TCRs. To determine the structure of Vγ9Vδ2 T-cell receptor, we generated two expression vectors containing all γδ TCR subunits linked by 2A ribosome skipping peptides (Fig. 1*A* and Fig. S1*A* and Table S1). A Twin-Strep tag and a FLAG tag were added to the C-termini of CD3ζ in tandem.

**Fig. 1.**
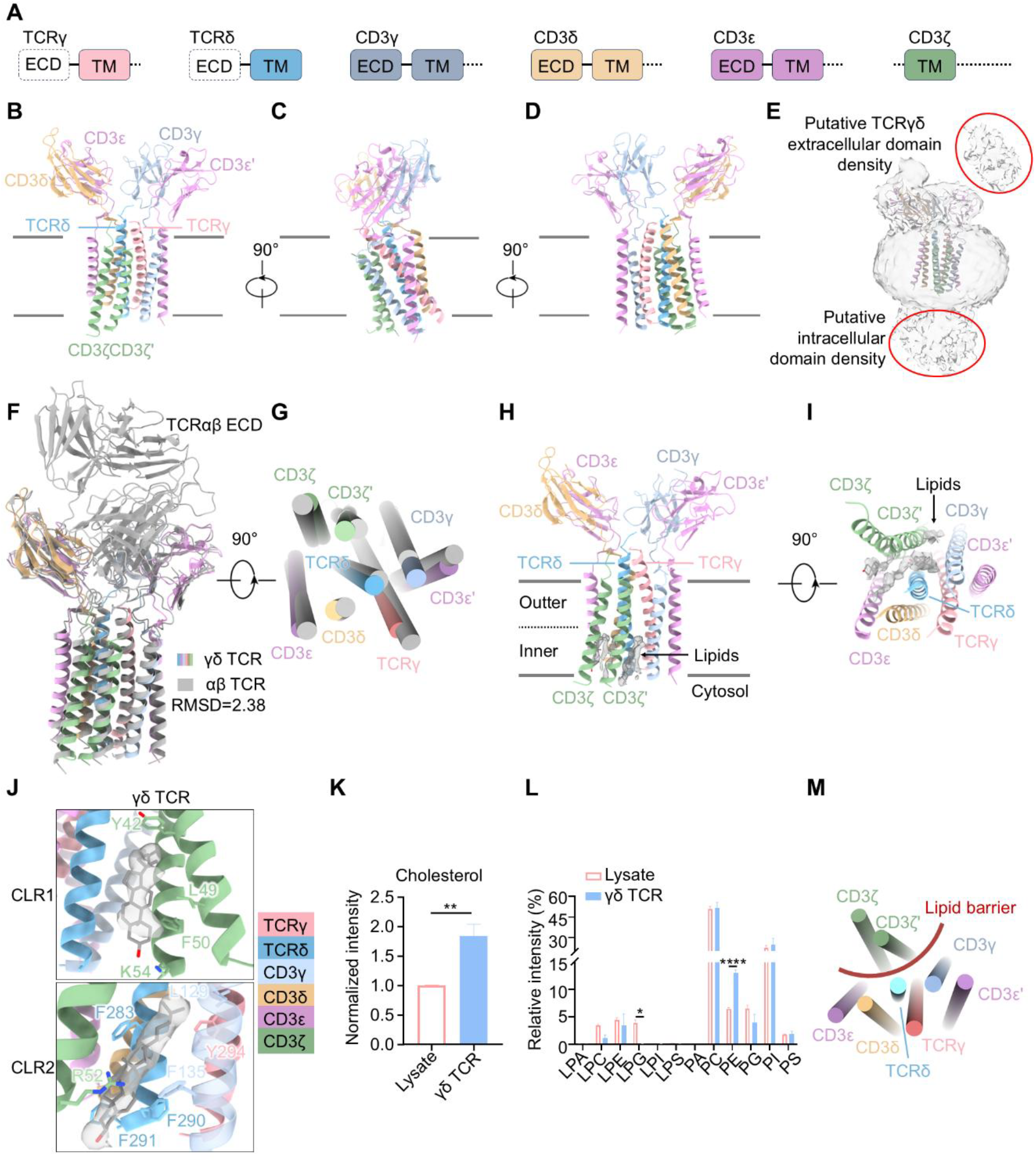
Cryo-EM structure and ordered lipids of γδ TCR. (***A***) Domain architectures for the γδ TCR. The dotted lines indicate flexible regions. (***B***-***D***) Ribbon diagrams of γδ TCR in three orthogonal views. Subunits are colored as follows: TCRγ (pink), TCRδ (light sky blue), CD3γ (light steel blue), CD3δ (burly wood), CD3ε (plum), and CD3ζ (dark sea green). Membrane boundaries are indicated by black lines. (***E***) Cryo-EM density of γδ TCR. (***F***) Overlaid ribbon diagram, showing the conformational changes of αβ TCR (PDB: 6JXR, gray) to γδ TCR (colored). (***G***) Comparison of the organization of the TM helices in the αβ TCR (gray) versus their counterparts in the γδ TCR (colored). (***H*** and ***I***) Ribbon diagrams of the overall structure of lipid-bound TM helices of TCR-CD3, shown in two orthogonal views. The surface indicates the EM densities of lipid molecules. Membrane boundaries are indicated by black lines. (***J***) Enlarged views of cholesterol (CLR) binding interfaces in the γδ TCR. (***K***) Quantification of cholesterol in γδ TCR-expressing Expi293F lysates or purified γδ TCR with untargeted lipidomics analysis (*n* = 3 biologically independent experiments; mean ± SD). Statistical significance was determined using two-tailed unpaired *t*-test. (***L***) Quantification of phospholipids in γδ TCR-expressing Expi293F lysates or purified γδ TCR with untargeted lipidomics analysis (*n* = 3 biologically independent experiments; mean ± SD). Statistical significance was determined using two-way ANOVA followed by Sidak’s multiple comparisons test. (***M***) Schematic diagram of how lipid molecules form a lipid barrier. Statistical significance: \**P* < 0.05; \*\**P* < 0.01; \*\*\*\**P* < 0.0001.

Surface expression of the γδ TCR on Expi293F cells was confirmed via flow-cytometry (Fig. S1*B* and C). The expressed γδ TCR complex was solubilized in glycol-diosgenin (GDN) micelles and purified by size exclusion chromatography (Fig. S1*D*). A monodispersed peak of the γδ TCR complex was collected, and validated by SDS-PAGE (polyacrylamide gel electrophoresis) and western blotting (Fig. S1*E* and *F*).

Next, we performed cryo-EM imaging of the purified complex and resolved the structure of the Vγ9Vδ2 TCR complex at an overall resolution of 3.3 Å (Fig. S2*A-G*). The γδ TCR adopts a Y-shaped architecture, with the extracellular domains (ECD) of CD3δε and CD3γε extending from the transmembrane region as two diverging arms at the top (Fig. 1*B-D*). The cryo-EM density unambiguously shows that the γδ TCR consists of one TCRγδ heterodimer, one CD3δε heterodimer, one CD3γε heterodimer, and one CD3ζζ homodimer (Fig. S2*E-G*). Notably, the extracellular domains of TCRγδ were not resolved in our cryo-EM map, except for some weak distal densities (Fig. 1*E*), suggesting substantial flexibility. This contrasts sharply with the more rigid ECD of TCRαβ in the αβ TCR (14-17) and is likely adopted to the antigen recognition and activation of the γδ TCR.

Superposition of our Vγ9Vδ2 TCR structure onto the published Vγ9Vδ2 TCR structure (PDB: 8JC0) (10) reveals a root-mean-square deviation (RMSD) of 1.180 Å, indicating no significant structural differences between the two models (Fig. S2*H*). Interestingly, the overall arrangement of the TCR and CD3 subunits in the γδ TCR is similar to that in the αβ TCR (RMSD = 2.38) (Fig. 1F), challenging previous speculation that the CD3 subunit geometry differs between the two receptors (6). Nevertheless, subtle differences were observed in the transmembrane region of the γδ and αβ TCRs (Fig. 1G), suggesting potential divergence in transmembrane signaling.

Together, our results reveal the structure of the Vγ9Vδ2 TCR and underscore the flexibility of the antigen-binding domains. This observation aligns with previous structural studies of the Vγ9Vδ2 TCR, and similar findings in other γδ TCR subtypes, including Vγ8Vδ3 and Vγ5Vδ1 TCRs (7-10), indicating that flexibility of the antigen-binding domains is a common feature of γδ TCR.

### A Lipid Barrier in the γδ TCR Complex

Previous studies have highlighted cholesterol as a key component of the TCR complexes (7-10, 18-20), but little is known about the phospholipids associated with the TCR complexes. In our γδ TCR structure, we observed multiple ordered lipid densities in the transmembrane region, including two cholesterols (CLR), three phospholipids (PL) in the inner leaflet of the γδ TCR (Fig. 1*H-J* and Fig. S2*I*). Two cholesterol-binding pockets, designated CLR1 and CLR2, were identified. CLR1 is formed primarily by residues Y42, L49, F50, and K54 of the CD3ζ. CLR2 involves a more extensive interface, comprising L129 and F135 from CD3γ; R52 from CD3ζ; Y294 from TCRγ; and F283, F290, and F291 from TCRδ (Fig. 1*J*).

To identify these bound lipids, we performed untargeted lipidomic analysis of the purified γδ TCR complex. Compared with whole-cell lysate, the γδ TCR complex showed marked enrichment of cholesterol and phosphatidylethanolamine (PE) species (Fig. 1*K* and *L*). Collectively, lipids are wedged between CD3ζζ and the rest of the γδ TCR complex, forming a continuous lipid “barrier” (Fig. 1*M*).

### Cross-linking of γδ TCR Complex by UCHT1 Fab

The anti-CD3ε antibody UCHT1 and its Fab fragment both enhance the tumor-killing activity of γδ T cells, serving as effective T cell agonists (19, 21, 22). To elucidate the mechanism by which UCHT1 Fab acts as a γδ T-cell agonist, we obtained UCHT1 IgG1 purified from the supernatant of UCHT1 monoclonal antibody hybridoma cells (Proteintech Group, Inc.) (23). We subsequently generated the UCHT1 Fab by ficin digestion followed by size exclusion chromatography purification (Fig. S3*A*). The purified UCHT1 Fab was analyzed by SDS-PAGE under reducing and non-reducing conditions and by blue native PAGE (Fig. 2*A*). Mass photometry and native mass spectrometry analysis indicate that the Fab fragment exists as a monomer in solution, with molecular weight of 52 ± 12.8 kDa and 48221 ± 0.1 Da, respectively (Fig. 2*B* and Fig. S3*B*). Electrophoretic mobility shift assay showed that the UCHT1 Fab binds to purified γδ TCR, resulting in shifts on blue native PAGE (Fig. 2C). Expi293F cells expressing Twin-Strep-mNeonGreen-tagged γδ TCR were treated with Fab, lysed, and subjected to Twin-Strep tag-based enrichment. Native PAGE analysis of the enriched samples revealed high-molecular-weight bands, indicating that UCHT1 Fab induces crosslinking of the γδ TCR on cell membranes (Fig. S3*C*). Consistently, cryo-EM imaging of the γδ TCR in complex with the Fab revealed net-like aggregates similar to those induced by UCHT1 IgG1, demonstrating that the UCHT1 Fab crosslinks the γδ TCR complex (Fig. 2*D* and Fig. S3*D*).

**Fig. 2.**
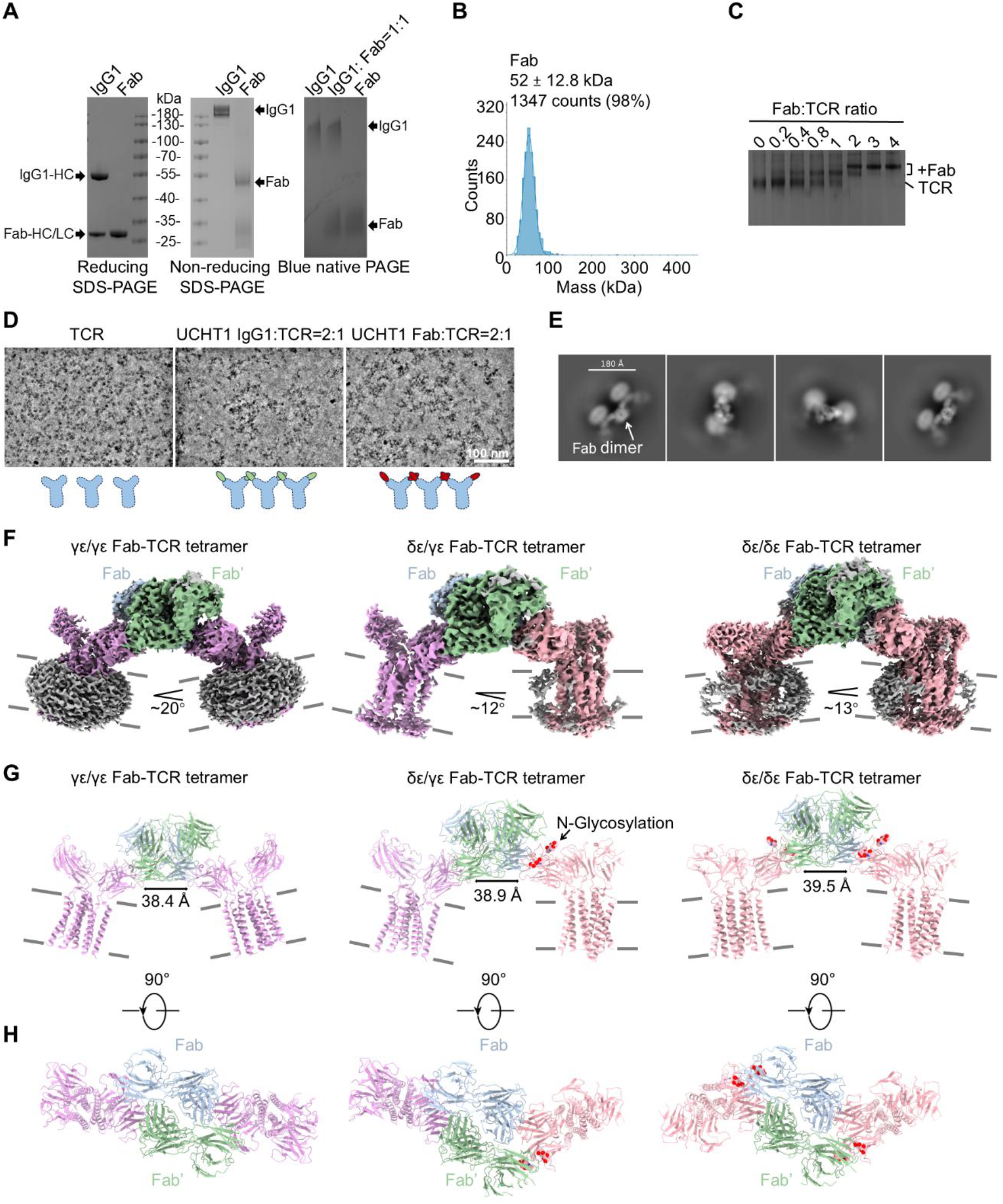
Cryo-EM structures of UCHT1 Fab-γδ TCR complexes. (***A***) Reducing SDS-PAGE (left), non-reducing SDS-PAGE (middle), and blue native PAGE (right) analyses of UCHT1 IgG1 and Fab. (***B***) Mass photometry of UCHT1 Fab. (***C***) Binding of Fab to γδ TCR assessed by blue native PAGE. The γδ TCR was mixed with Fab at the indicated molar ratios prior to loading on the gel. (***D***) Representative low-pass filtered (to 14 Å) cryo-EM micrographs of the γδ TCR, UCHT1 IgG1-γδ TCR complex, and Fab-γδ TCR complex, which were imaged on a FEI Titan Krios G4 microscope operated at 300 kV. The schematic diagrams below illustrate the complex configurations. (***E***) 2D class averages of Fab-γδ TCR complexes. (***F***) Cryo-EM densities of γε/γε Fab-TCR tetramer (left), δε/γε Fab-TCR tetramer (middle), and δε/δε Fab-TCR tetramer (right) shown in side view. The γδ TCRs which bind to Fab through CD3γε or CD3δε are colored in plum and pink, respectively. The Fabs are colored in blue and green, respectively. Membrane boundaries are indicated by black lines. (***G*** and ***H***) Ribbon diagrams of γε/γε Fab-TCR tetramer (left), δε/γε Fab-TCR tetramer (middle), and δε/δε Fab-TCR tetramer (right) are shown in two orthogonal views. The sphere models indicate N-glycosylation sites and colored by element.

To investigate the mechanism of γδ TCR crosslinking by the UCHT1 Fab fragment, we incubated the Fab with the γδ TCR, and performed cryo-EM study (Fig. S4). Interestingly, most of the 2D classes revealed complexes consisting of two Fab fragments and two TCR complexes (Fig. 2*E*). 3D reconstruction of the Fab-TCR tetramers (Fig. S4) showed that the UCHT1 Fab dimerizes to bridge two TCR complexes. The cryo-EM densities at the TCR region are ambiguous, though the local resolution of the Fab dimer region is high. To address this, we performed local 3D classification focused on the TCR region and found that the TCR binds to Fab either through CD3γε or CD3δε (Fig. S4). Accordingly, we classified all particles into three groups based on their Fab-TCR binding sites (γε/γε, δε/γε, and δε/δε) and performed 3D reconstructions separately (Fig. S4). This yielded three cryo-EM structures of the Fab-TCR tetramers: γε/γε, δε/γε, and δε/δε, with resolutions of 3.29 Å, 3.35Å, 3.08 Å, respectively (Fig. 2*F-H* and Fig. S5).

Notably, the three Fab-TCR tetramers exhibit distinct conformational distributions, with 49.2%, 31.4%, and 19.4% of the particles adopting the δε/γε, δε/δε, and γε/γε conformations, respectively (Fig. S4). Within these Fab-TCR tetramers, we observed noticeable curvature (Fig. 2*F*), suggests that Fab binding on the plasma membrane may generate mechanical forces.

### Unexpected Dimerization of UCHT1 Fab

To elucidate the structural basis of the UCHT1 Fab dimerization, we first examined the dimerized Fab fragments and identified three dimer interfaces, all mediated by the variable domains of the Fab (Fig. 3*A*). The first binding interface consists of two identical β-strands forming an anti-parallel β-sheet at the C2 symmetry axis (residues T_7_TSS_10_; Fig. 3*B*). The other two interfaces are symmetrical and contain a network of hydrogen bonds involving the N-terminus of the light chain (residues D1, Q3, and S26), and residues H41, N44, and E89 from the heavy chain (Fig. 3*C*). In mass photometry, size exclusion chromatography, and native mass spectrometry, free UCHT1 Fab eluted as a monomer (Fig. 2*B* and Fig. S3*A* and *B*), indicating that the dimerization of the Fab is induced by the TCR binding. In addition to the interaction between the UCHT1 Fab variable domain and CD3ε, we observed that the constant domain of the Fab light chain is in close proximity to the CD3δ ECD. Upon Fab binding, the CD3δ ECD undergoes conformational changes, with two N-glycosylation sites becoming ordered (Fig. 3*D* and *E*), suggesting potential interactions between the Fab constant domain and the CD3δ ECD. Thus, in addition to Fab-Fab interactions, Fab-TCR interactions may also contribute to the dimerization of UCHT1 Fab.

**Fig. 3.**
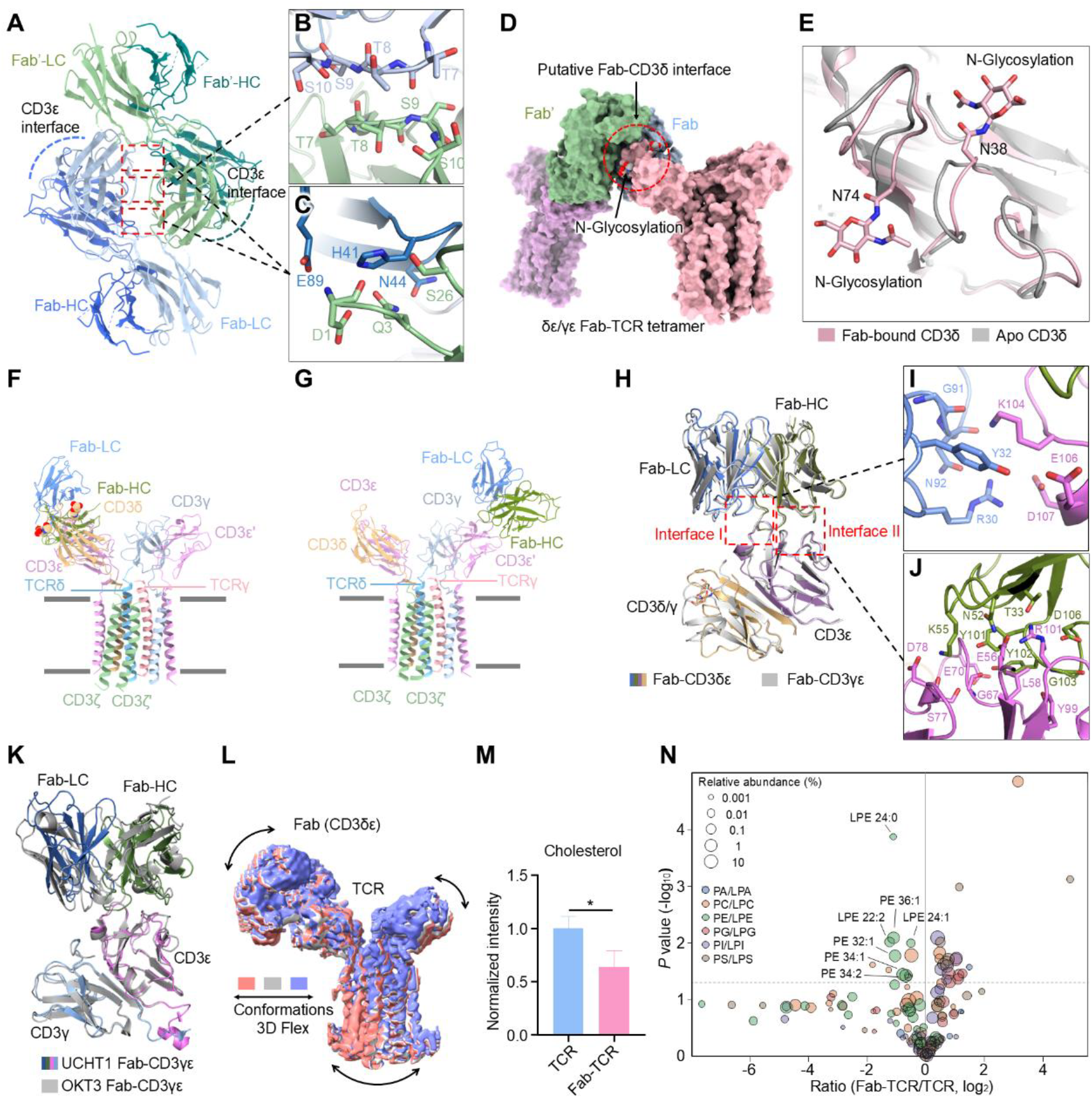
UCHT1 Fab dimer and Fab-TCR complex. (***A***) Ribbon diagram of Fab-Fab dimer. One Fab heavy and light chains are colored in royal blue and light steel blue respectively, and another one Fab heavy and light chains are colored in teal and dark sea green respectively. Two dotted lines indicate the interfaces between the variable domain of Fab heavy chain and CD3ε. Red boxes indicate the interfaces between Fab-Fab dimer. (***B*** and ***C***) Enlarged views of the Fab-Fab dimer interfaces. (***D***) Cryo-EM surface map of δε/γε Fab-TCR tetramer. Red circle indicates putative Fab-CD3δ interface. Two sphere models indicate N-glycosylation sites and colored by element. (***E***) Enlarged view of overlaid ribbon diagram, showing the conformational changes of apo γδ TCR (gray) to Fab-bound γδ TCR (pink). (***F*** and ***G***) Ribbon diagrams of Fab-CD3δε-TCR and Fab-CD3γε-TCR, respectively. Fab heavy and light chains are colored in green and blue respectively. (***H***) Enlarged view of overlaid ribbon diagram, Fab-CD3γε-TCR is gray and Fab-CD3δε-TCR is colored. Red boxes indicate two interfaces between Fab and CD3ε. (***I***) Enlarged view of Interface I between Fab light chain (blue) and CD3ε (purple). (***J***) Enlarged view of Interface II between Fab heavy chain (green) and CD3ε (purple). (***K***) Enlarged view of overlaid ribbon diagram, OKT3 Fab-CD3γε-TCR (PDB: 1SY6) is gray and UCHT1 Fab-CD3γε-TCR is colored. (***L***) Three overlaid convected densities from the 3D Flex model of the Fab-CD3γδ-TCR. (***M***) Quantification of cholesterol in purified γδ TCR or Fab-γδ TCR with untargeted lipidomics (*n* = 3 biologically independent experiments; mean ± SD). Statistical significance was determined using two-tailed unpaired *t*-test. Statistical significance: \**P* < 0.05. (***N***) Vocanol plot of quantified phospholipids in purified γδ TCR or Fab-γδ TCR with untargeted lipidomics (*n* = 3 biologically independent experiments). Statistical significance was determined using two-sided unpaired Student’s *t*-tests. The significantly decreased PE/LPE species in Fab-γδ TCR samples are labelled accordingly.

### Interactions Between Fab and CD3ε of The γδ TCR Complex

To improve the resolution of the TCR regions in the Fab-TCR tetramer, we performed local refinement of the Fab-CD3δε-TCR and Fab-CD3γε-TCR regions, yielding two structures with resolutions of 3.24 Å and 3.69 Å, respectively (Fig. 3*F* and *G* and Fig. S6*A* and *B*). In both Fab-TCR complex structures, the binding interface between the variable domains of the Fab and the ECD of the CD3ε chain are nearly identical (Fig. 3*H*). This binding interface involves key residues Y32, R30, G91, and N92 from the Fab light chain, and residues K104, E106, and D107 from CD3ε (Fig. 3*H* and *I*). The variable domain of the Fab heavy chain engages more extensive interactions with CD3ε, involving residues T33, N52, K55, Y101, Y102, G103, and D106 from the Fab heavy chain and residues E56, L58, G67, E70, S77, D78, and Y99 from CD3ε (Fig. 3*H* and *J*).

Although both OKT3 and UCHT1 bind CD3ε, only UCHT1 robustly induces γδ TCR-proximal intracellular signaling, including Ca^2+^influx and phosphorylation of downstream signaling proteins, and greatly enhances tumor cell killing by γδ T cells (21, 22). To compare their binding to CD3ε, we overlaid the OKT3 Fab-TCR complex with the UCHT1 Fab-TCR complex (Fig. 3*K*). Surprisingly, OKT3 and UCHT1 bind to CD3ε in a similar manner, suggesting that their functional differences may result from Fab dimerization rather than their binding interfaces to CD3ε.

### Conformation Changes of Fab-TCR

To investigate whether Fab binding induces conformational changes in the TCR complex, we overlaid the local-refined Fab-CD3δε-TCR and Fab-CD3γε-TCR structures with the apo TCR, yielding RMSD values of 1.326 Å and 1.205 Å, respectively (Fig. S6*C* and *D*). Superposition of our Vγ9Vδ2 TCR-UCHT1 Fab structure with the published Vγ8Vδ3 TCR-UCHT1 Fab (PDB: 9CI8) (9) and Vγ9Vδ2 TCR-OKT3 Fab (PDB: 9CQ4) (7) complexes yielded RMSD values of 1.115 Å and 1.200 Å, respectively (Fig. S6E and *F*). These results support a conserved overall architecture of γδ TCR complexes despite differences in γδ TCR subtype or bound Fabs.

Although the overall architecture of apo and Fab-TCR complexes is similar, we observed subtle yet noticeable changes in the extracellular and transmembrane regions of CD3 subunits, indicating conformational changes upon Fab binding. 3D variability analysis of the Fab-CD3δε-TCR revealed continuous motion in the extracellular domains and transmembrane regions of CD3 dimers. These conformation changes were further confirmed by the 3D Flex analysis in CryoSPARC (24). The overlaid convected densities from the 3D Flex model suggest motion involving the closure of the extracellular domains of CD3δε and CD3γε (Fig. 3*L*), which propagates to the transmembrane region, potentially impacting the cytosolic side. These results support the idea that the CD3 dimers serve as mechano-sensors capable of transducing forces into intracellular signals (25).

To further characterize the TCR-associated lipids upon Fab stimulation, we performed untargeted lipidomic analysis of the Fab-γδ TCR complex. Our data revealed a marked reduction of cholesterol and several phosphatidylethanolamine (PE) species in the Fab-γδ TCR complex relative to the unstimulated γδ TCR complex (Fig. 3*M* and *N*). Consistently, the Fab-TCR complex retains a well-resolved CLR2 density, in contrast to the diffuse densities of CLR1 and phospholipids (Fig. S6*G*). Together, these observations suggest that Fab binding may influence the local lipid environment of the γδ TCR complex.

### Fab Dimerization Enhances TCR Activation

To systematically evaluate the role of Fab dimerization in γδ TCR activation, we employed recombinant expression to generate recombinant Fab proteins (rFab) and their mutants. The UCHT1 Fab sequence was determined by integrating the previously reported sequence (9), *de novo* sequencing of the UCHT1 IgG1, N-terminal Edman sequencing of the UCHT1 light chain, and the cryo-EM structure of the UCHT1 Fab (Tables S1 and S2 and Fig. S7*A* and Fig. 2*F*). The OKT3 Fab sequence was derived from the reported OKT3 Fab sequence (26, 27). We included an rFab-L70F mutant (Fig. S7*B*) to account for a single-residue difference (L70 versus F70 in the heavy chain) between our UCHT1 Fab and the published sequence (9). UCHT1 and OKT3 rFabs were purified from the culture supernatant using Ni-NTA affinity chromatography followed by size-exclusion chromatography (Fig. 4*A* and Fig. S7*C* and *D*). Mass photometry demonstrated that all rFabs are predominantly monomeric (Fig. 4*B* and Fig. S7*B* and *D*).

**Fig. 4.**
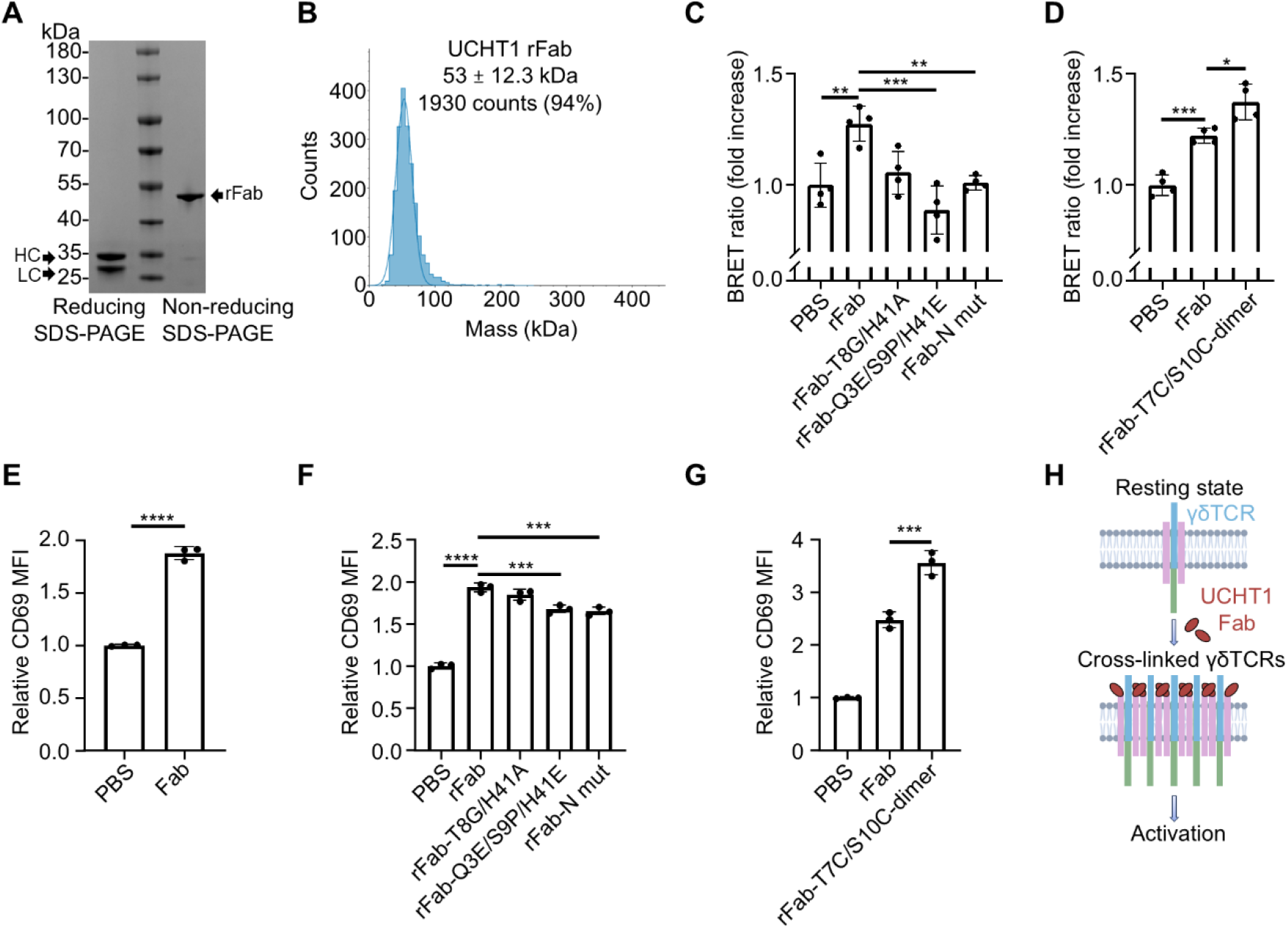
Fab dimerization enhances TCR activation. (***A***) Reducing and non-reducing SDS-PAGE analyses of UCHT1 recombinant Fab (rFab). (***B***) Mass photometry of UCHT1 rFab. (***C*** and ***D***) BRET ratios in Expi293F cells co-expressing γδ TCR-NanoLuc and γδ TCR-mNeonGreen upon treatment with PBS (vehicle control), rFab, or rFab mutants (*n* = 4 biologically independent experiments; mean ± SD). Dimer-disrupting mutants: rFab-T8G/H41A (T8G in the light chain and H41A in the heavy chain), rFab-Q3E/S9P/H41E (Q3E/S9P in the light chain and H41E in the heavy chain), and rFab-N mut (deleting residues D_1_IQ_3_ and replacing residues T_7_TSS_10_ with GSGS in the light chain). Dimer-stabilizing mutant: rFab-T7C/S10C (T7C/S10C in the light chain). (***E***-***G***) γδ TCR-expressing Jurkat cells were stimulated with PBS, ficin-generated Fab (***E***), rFab, or rFab mutants (***F*** and ***G***) for 3 h, followed by staining with PE-conjugated anti-human CD69 antibody for flow cytometric analysis of CD69 upregulation (*n* = 3 biologically independent experiments; mean ± SD). MFI, median fluorescence intensity. (***H***) Schematic illustration of γδ TCR clustering as a general mechanism leading to γδ TCR activation. *P* values were calculated using one-way ANOVA followed by Tukey’s multiple comparisons test (***C, D, F***, and ***G***) or by two-tailed unpaired *t*-test (***E***). Statistical significance: \**P* < 0.05; \*\**P* < 0.01; \*\*\**P* < 0.001; \*\*\*\**P* < 0.0001.

To elucidate the function of the dimerized UCHT1 Fab, we designed mutations to either disrupt or stabilize the dimer. Structural analysis identified three Fab dimer interfaces: Interface 1, located at the light-chain segment T_7_TSS_10_, and two additional hydrogen-bonding interfaces involving light-chain residues D1, Q3, and S26 with heavy-chain residues H41, N44, and E89 (Fig. 3*A*-*C*). Based on these interfaces, we purified a series of dimer-disrupting mutants, including rFab-T8G/H41A (T8G in the light chain and H41A in the heavy chain to disrupt key hydrogen bonds), rFab-Q3E/S9P/H41E (Q3E/S9P in the light chain and H41E in the heavy chain to introduce charge repulsion and β-sheet destabilization), and rFab-N mut (deleting residues D_1_IQ_3_ and replacing residues T_7_TSS_10_ with GSGS in the light chain to disrupt all interfaces) (Fig. S8). In parallel, rFab-T7C/S10C (T7C/S10C in the light chain) was designed to stabilize Fab dimerization through an intermolecular disulfide bond (Fig. S9). Mass photometry confirmed that all variants remained monomeric except for rFab-T7C/S10C-dimer, which formed stable dimers (Figs. S8 and S9).

To evaluate the impact of Fab dimerization on binding affinity, we generated Jurkat cells expressing γδ TCR instead of αβ TCR (Fig. S10*A*-*C*) and assessed rFabs binding via flow cytometry. The OKT3 rFab exhibited a markedly higher K_D_ than the UCHT1 rFab, despite their similar CD3ε-binding modes (7, 26) (Fig. S10*D* and Fig 3*K*). The dimer-disrupting mutants of the UCHT1 rFab showed increased K_D_ values, particularly rFab-Q3E/S9P/H41E (Fig. S10*D*). These findings suggest that dimerization enhances the binding of UCHT1 rFab.

To determine whether UCHT1 Fab induces γδ TCR crosslinking on cells, we performed bioluminescence resonance energy transfer (BRET) assays. Structural analysis of the Fab-γδ TCR complexes showed that the distances between the two TCR molecules ranged from 38.4 to 39.5 Å (Fig. 2*G*), which falls within the effective detection range of the NanoLuc-mNeonGreen BRET pair used in our assay, based on its 51 Å Förster radius (R_0_) (28). We showed that UCHT1 rFab induced the close proximity between Vγ9Vδ2 TCR molecules while all dimer-disrupting mutations reduced this effect, particularly rFab-Q3E/S9P/H41E and rFab-N mut (Fig. 4*C*). The rFab-T7C/S10C-dimer further enhanced TCR clustering (Fig. 4*D*). Thus, rFab dimerization drives TCR crosslinking.

Next, we assessed the downstream signaling of the γδ TCR. The UCHT1 IgG1 efficiently induced CD69 upregulation in γδ TCR-Jurkat cells (Fig. S10*E*), validating the robustness of our assays. UCHT1 Fab generated by ficin digestion also promoted CD69 upregulation (Fig. 4 *E*). Recombinant UCHT1 Fab exhibited comparable activity, confirming that it is fully functional (Fig. S10*F*). Moreover, the rFab-L70F (PDB: 9CI8) (9) exhibited similar potency to our rFab, indicating that our rFab is functional equivalent to the previously reported rFab (9) (Fig. S10*F*).

Having validated UCHT1 rFab-induced γδ TCR activation, we proceeded to investigate dimer-disrupting and dimer-stabilizing rFab mutants. All dimer-disrupting mutants exhibited impaired activity, with rFab-Q3E/S9P/H41E and rFab-N mut showing the strongest inhibition (Fig. 4*F*). Conversely, dimer-stabilizing variant rFab-T7C/S10C significantly enhanced CD69 expression relative to rFab, suggesting that enforced Fab dimerization promotes more effective TCR activation than spontaneous dimer formation of rFab (Fig. 4*G*). Compared with rFab, OKT3 rFab exhibited a lower BRET signal and a reduced capacity to induce CD69 upregulation. (Fig. S10*G* and *H*). Taken together, these results demonstrate that Fab dimerization enhances γδ TCR activation.

## Discussion

We determined structures of the γδ TCR in both its apo and Fab-bound states. Although the overall architecture of the γδ TCR closely resembles that of the αβ TCR, a striking distinction is the extensively flexibility of TCRγδ extracellular domains, which is consistent with previous studies (7-10). This difference in flexibility likely reflects the distinct ligand-recognition strategies of these two receptor classes, as γδ TCRs engage with structurally diverse ligands rather than the MHC-dependent ligands of αβ TCRs.

In our 3.3 Å γδ TCR structure, we identified a set of ordered lipids, including phospholipids and cholesterols, that form a “barrier” in the inner leaflet, spatially separating the CD3ζζ dimer from other subunits of the γδ TCR. Cholesterol has been shown to inhibit γδ TCR activation (10). This lipid-mediated separation of CD3ζζ likely prevents TCR self-activation, although the exact mechanism remains unclear.

Our structure of the UCHT1 Fab-γδ TCR complex reveals an unexpected spontaneous dimerization. Further analysis suggested that free UCHT1 Fab is monomeric and dimerizes only upon binding to the γδ TCR. This observation explains why UCHT1 and UCHT1 Fab can promote γδ TCR-proximal intracellular signaling, including Ca^2+^ influx and the phosphorylation of downstream signaling proteins, and greatly enhances tumor cell killing by γδ T cells, whereas other TCR-binding antibodies such as OKT3 can not (21, 22). Although UCHT1 and OKT3 rFabs bind to CD3ε in a similar manner, it is plausible that the increased multivalency from Fab dimerization likely accounts for γδ TCR activation. This is supported by a previous study showing that increasing the multivalency of OKT3 with a secondary antibody activates the γδ TCR (21).

Importantly, BTN receptor complexes have been identified as physiological ligands of the Vγ9Vδ2 TCR (29-33). Two recent study have shown that the HMBPP-BTN2A1-BTN3A1-BTN3A2 natural ligand complex provides two spatially distinct binding sites for the γδ TCR, enabling simultaneous engagement of two γδ TCRs and promoting γδ TCR crosslinking and activation (8, 34). These findings provide evidence that multivalent engagement of the γδ TCR extends to the context of natural ligands. Together, natural (BTN-mediated) and engineered (Fab-mediated) γδ TCR clustering supports a general role of ligand valency in receptor activation.

Previous structural studies of Vγ9Vδ2 TCR complexes with anti-TCRδ or anti-CD3ε (OKT3) Fab did not show receptor crosslinking (7), indicating that not all γδ TCR antibodies are capable of driving receptor clustering. A recent study proposed that the αβ TCR activates through two distinct mechanisms, digital and analog activation, whereas the γδ TCR relies exclusively on analog activation (35). The analog activation requires ligand-induced TCR clustering. Consistent with this principal, dimerization of the UCHT1 Fab may increase its multivalency and thereby facilitate γδ TCR clustering. Therefore, our findings and the recent studies (8, 34) underscore multivalency as a key factor for antibody-induced activation of the γδ TCR and suggest that clustering of the γδ TCR, whether induced by antibody/Fab crosslinking or ligand-binding, represents a general mechanism of γδ TCR activation (Fig. 4*H*). The unique dimerization mode of the UCHT1 Fab provides a foundation for the future development of dimerized Fab as therapeutic agents.

## Materials and Methods

### Cell Lines

Expi293F cells (Thermo Fisher Scientific) were cultured in SMM 293T-II medium (Sino Biological) at 37 °C under 5% CO_2_, 110 rpm. HEK293T cells (Procell) and Jurkat T cells (clone E6-1; Procell) were maintained at 37 °C in a humidified atmosphere containing 5% CO_2_ in DMEM and RPMI 1640 medium (Gibco), respectively, supplemented with 10% (v/v) fetal bovine serum (FBS).

### γδ TCR Protein Expression and Purification

We selected a prototypical Vγ9Vδ2 TCR derived from G115 clonotypes (13). The full-length TCRγ and TCRδ coding sequences were subcloned into pcDNA3.1 vector, separated by a T2A peptide. The full-length sequences of CD3 subunits were designed as follows: CD3ε-T2A-CD3γ-P2A-CD3δ-E2A-CD3ζ-Twin-Strep tag-FLAG tag, and subcloned into the pcDNA3.1 vector.

When the cell density reached 3×10^6^ cells /ml, 0.5 mg each of the TCR-pcDNA3.1 and CD3-pcDNA3.1 plasmids and 5 mg of PEI (25 kDa, Polysciences) were mixed in 50 mL of fresh medium and incubated for 30 min before transfecting 1 L of Expi293F cells. Transfected cells were incubated at 37 °C and 5% CO_2_ for 24 h, then 10 mM sodium butyrate was added and incubated for an additional 48 h. Surface expression of the γδ TCR-CD3 complex was confirmed by flow cytometry. Cells were harvested by centrifugation, washed with PBS, and stored at -80 °C.

Cells were thawed and resuspended in buffer containing 25 mM HEPES, pH 8.0, 150 mM NaCl, 1% n-dodecyl-β-D-maltopyranoside (DDM, Anatrace), 0.1% cholesteryl hemisuccinate tris salt (CHS, Anatrace), and SIGMAFAST™ Protease Inhibitor Cocktail (Sigma-Aldrich). The mixture was stirred at 4 °C for 4 h and then clarified by centrifugation at 18,000 rpm for 30 min. Then the supernatant was incubated with Strep-Tactin resin (IBA) for 2 h at 4 °C. The resin was collected and washed with 10 column volumes of wash buffer containing 25 mM HEPES, pH 8.0, 300 mM NaCl, and 0.02% GDN (Anatrace). The γδ TCR complex was eluted with wash buffer supplemented with 4 mM desthiobiotin, concentrated to 1 mL in a 100 kDa MWCO Amicon Ultra centrifugal filter, and further purified by size-exclusion chromatography (Superose 6 increase 10/300 GL, GE Healthcare) in running buffer (25 mM HEPES, pH 8.0, 150 mM NaCl, 0.02% GDN). Purity of the proteins was monitored using SDS-PAGE under reducing condition and visualized by Coomassie blue staining (Lablead). The size-exclusion chromatography fractions were analysed by western blot using anti-FLAG antibodies (Proteintech Group, Inc.) at a dilution of 1:5,000, and anti-TCRδ antibodies (Santa Cruz Biotechnology) at a dilution of 1:500. The secondary goat anti-mouse antibody (Proteintech Group, Inc.) was used at a dilution of 1:10,000.

### Flow Cytometry Analysis

Cells were washed with FPBS (1% FBS in PBS) and stained for flow cytometry with APC-conjugated anti-human TCR-Vδ2 antibody (clone B6, BioLegend, diluted 1:25), PE-conjugated anti-human CD3ε antibody (clone UCHT1, BioLegend, diluted 1:25), or BV421-conjugated anti-human CD3ε antibody (clone OKT3, BioLegend, diluted 1:25) for 30 min at 4 °C in the dark. Cells were subsequently washed with FPBS. Data were acquired on a CytoFlex (Beckman) and analyzed with FlowJo software (FlowJo LLC). Cells were gated by FSC-A versus SSC-A to exclude debris and then by FSC-A versus FSC-H to exclude adhered cells (Fig. S1*C*).

### Preparation of Fab Fragments by Ficin Digestion

For preparation of Fab Fragments, ficin digestion method was adapted from previous methods (22, 36). UCHT1 IgG1 (Proteintech Group, Inc.) was dialyzed overnight at 4 °C against 1 L of buffer containing 50 mM Tris-HCl (pH 7.0) and 2 mM EDTA. Ficin (Sigma-Aldrich) was mixed with UCHT1 IgG1 at a 1:30 weight ratio in cutting buffer (50 mM Tris-HCl, pH 7.0, 2 mM EDTA, 25 mM cysteine) and incubated in a 37 °C water bath with gentle shaking. The reaction was terminated by addition of 30 mM N-ethylmaleimide (Aladdin). The digestion mixture was dialyzed against buffer containing 50 mM HEPES (pH 7.0) and 50 mM NaCl and passed through a HiTrap Capto S column (Cytiva) to remove residual ficin(37). The sample was diluted in buffer containing 150 mM NaCl and 20 mM Na_2_HPO_4_ (pH 7.2) before loading onto a HiTrap MabSelect PrismA column (Cytiva). The flow-through fractions were collected and concentrated in a 10 kDa MWCO Amicon Ultra centrifugal filter, then injected into a Superdex 200 Increase 10/300 GL (Cytiva) equilibrated to size-exclusion chromatography buffer (25 mM HEPES, pH 8.0, 150 mM NaCl). Protein purity was assessed by reducing and non-reducing SDS-PAGE and blue native PAGE, followed by Coomassie blue staining. A protein standard mixture (Sigma-Aldrich) was analyzed on the same column under identical buffer conditions. In cellular functional assays, the Fab fragment was buffer-exchanged into PBS using a 10 kDa MWCO filter (Amicon).

### Cryo-EM Data Collection and Image Processing

Cryo-EM grids were prepared using the γδ TCR at a concentration of ∼2 mg/mL. Purified γδ TCR was mixed with UCHT1 Fab (1:2 molar ratio) or UCHT1 IgG1 (1:2 molar ratio), and incubated for 30 min on ice. UltrAuFoil 1.2/1.3 grids were used for the γδ TCR sample and UCHT1 IgG1-γδ TCR complex. Quantifoil R1.2/1.3 grids were used for the Fab-γδ TCR complex. In each case, A 3-μL aliquot of the sample was applied to a glow-discharged grid mounted in the chamber of the Vitrobot Mark IV (Thermo Fisher Scientific) at 8 °C and 100% humidity. After blotting for 3.5 s, the grid was quickly plunged into liquid ethane cooled by liquid nitrogen. These grids were screened on a FEI Glacios microscope (Core Facility of Wuhan University) operated at 200 kV and equipped with a Ceta D CMOS camera. Grids with optimal ice thicknesses and particle densities were chosen for cryo-EM data collection. All data collection was performed using a FEI Titan Krios G4 microscope (Core Facility of Wuhan University) operated at 300 kV and equipped with a Gatan K3 direct electron detector. For the γδ TCR, 7,028 movies were collected using Thermo Scientific EPU software in counting mode, with each movie containing 40 total frames, an accumulated dose of 50 electrons per Å^2^, and a pixel size of 0.84 Å. For the Fab-γδ TCR complex and the UCHT1 IgG1-γδ TCR complex, 7,811 and 500 movies were collected, respectively, using Thermo Scientific EPU software in counting mode. Each movie comprised 40 total frames, with an accumulated dose of 50 electrons per Å^2^ and a pixel size of 0.824 Å. All datasets were processed in CryoSPARC (24).

For the γδ TCR, 5,548 images were retained after patch-based motion correction and CTF estimation, from which 7,322,054 particles were picked using autopick and handpick. After 2 rounds of 2D classification, 554,539 particles were selected for ab-initio reconstruction followed by heterogeneous refinement with C1 symmetry. The particles in class 1 were selected to do non-uniform refinement and local refinement with C1 symmetry. Particles were used in one additional round of ab-initio reconstruction followed by heterogeneous refinement with C1 symmetry. The best class, with 146,233 particles, was used for non-uniform refinement and local refinement with C1 symmetry, and followed by local resolution estimation and local filtering. The resulting map has a resolution of 3.31 Å.

For the Fab-γδ TCR complex, 7,577 images were retained for patch-based motion correction and CTF estimation, from which 3,738,972 particles were extracted using autopick and template picker. After 2 rounds of 2D classification, 443,947 particles were selected for ab-initio reconstruction, followed by heterogeneous refinement with C1 symmetry. The best class was used for ab-initio reconstruction, followed by non-uniform refinement with C2 symmetry, which resulted in an initial volume with a 2.82 Å resolution. This volume was subjected to symmetry expansion with C2 symmetry, followed by focused 3D classification, producing two classes. Class 0 was subjected to local refinement with C1 symmetry, followed by 3D flex reconstruction and sharpening tools, resulting in a Fab-CD3δε-TCR map with a resolution of 3.24 Å. Class 1 was subjected to local refinement with C1 symmetry, followed by 3D classification and local refinement with C1 symmetry, resulting in a Fab-CD3γε-TCR map with a resolution of 3.69 Å. We designated the tetramers with TCR(CD3γε)-Fab from the two conformations as set A, and the tetramers with TCR(CD3δε)-Fab as set B. Subtracting A from B, we performed non-uniform refinement with C2 symmetry, generating a δε/δε Fab-TCR map with a resolution of 3.08 Å. Subtracting B from A, we performed non-uniform refinement with C2 symmetry, generating a γε/γε Fab-TCR volume. This volume was then intersected with the Fab-CD3γε-TCR, followed by non-uniform refinement with C2 symmetry, resulting in a γε/γε Fab-TCR map with a resolution of 3.29 Å. Intersecting A with B, and followed by non-uniform refinement with C1 symmetry, generating a δε/γε Fab-TCR volume. This volume was then intersected with the Fab-CD3γε-TCR, followed by non-uniform refinement with C1 symmetry, resulting in a δε/γε Fab-TCR map with a resolution of 3.35 Å.

### Model Building and Structure Refinement

6JXR was used as the initial model for building the γδ TCR, and then the γδ TCR was used to build the Fab-γδ TCR complex. Model building was performed in Coot (38), and real-space refinement was carried out in PHENIX (39). Model quality was assessed using MolProbity (40). Statistical details can be found in Table S3. Figures were prepared in PyMOL and UCSF ChimeraX (41).

### Untargeted Lipidomics Analysis

The γδ TCR-expressing Expi293F cells were resuspended in buffer containing 25 mM HEPES (pH 8.0), 150 mM NaCl, 1% DDM, 0.1% CHS, and SIGMAFAST™ Protease Inhibitor Cocktail (Sigma-Aldrich). The cell suspension was stirred at 4 °C for 4 h and then clarified by centrifugation at 18,000 rpm for 30 min. The supernatant was filtered through a 0.45-μm syringe filter and used as the cell lysate. Purified γδ TCR was mixed with UCHT1 Fab (1:2 molar ratio) incubated for 30 min on ice. The protein sample was desalted using a Micro Bio-Spin P-6 Gel Column (Bio-Rad Laboratories. Inc) to remove unbound lipids. Lipids from recombinant proteins and cell lysate were subsequently extracted using the methyl tert-butyl ether (MTBE)/methanol method (42). Briefly, 40 μL of each sample was mixed with 300 μL methanol and 1 mL MTBE, and then incubated at room temperature for 10 min. Following the addition of 250 μL H_2_O, phase-separation was induced by centrifugation at 400g for 10 min. The upper (organic) phase was transferred to a new glass vial. The lower (aqueous) phase was re-extracted with 400 μL of a solvent mixture (MTBE/methanol/H_2_O:10/3/2.5). The organic phase was combined and dried with nitrogen flow in a fume hood.

The dried lipids were reconstituted in 60% acetonitrile (ACN) for subsequent LC-MS/MS analysis. The lipids were separated on a 15cm C18 column using a Vanquish Neo LC System. Separation was performed at 50 °C using a binary buffer system and a gradient of 20% to 99% buffer B at a flow rate of 300 nL/min over 20 min. (Buffer A of 60% ACN, 40% H2O, 10 mM ammonium formate, 0.1% formic acid and buffer B of 90% isopropanol: 10% ACN, 10 mM ammonium formate, 0.1% formic acid). The eluted lipids were delivered via a nanospray source to an Orbitrap Ascend Tribrid mass spectrometer (Thermo Scientific). Typical MS settings were a spray voltage of 1.8 kV, capillary temperature of 300 °C, and RF lens of 40%. The MS instrument was operated in small molecule mode with data-dependent acquisition. Full-scan MS spectra were acquired in the Orbitrap (m/z 300-2,000) at a resolution of 120,000. HCD fragmentation was performed with stepped collision energies of 25% and 30% and AGC target of 150%. The lipidomics raw data were processed using MS-DIAL (v5.5) software. Only peak features with reference-matched MS/MS spectra were selected for lipid quantification. All lipid identification and quantification were manually checked and normalized.

### Co-immunoprecipitation Assay

Vδ2-E2A-Vγ9 TCR-mNeonGreen-pcDNA3.1 and CD3-Twin-Strep tag-FLAG tag-pcDNA3.1 plasmids were transfected into 6×10^7^ Expi293F cells. After 72 h, cells were resuspended in 500 μL PBS, and 0, 30, or 60 μg Fab was added followed by incubation at 4 °C for 1 h. Cells were then lysed in lysis buffer containing 25 mM HEPES (pH 8.0), 150 mM NaCl, 1% DDM, 0.1% CHS, and SIGMAFAST™ Protease Inhibitor Cocktail. Cleared supernatants were incubated with Strep-Tactin resin (IBA) for 2 h at 4 °C. The resin was collected and washed with 10 column volumes of wash buffer containing 25 mM HEPES (pH 8.0), 300 mM NaCl, and 0.02% GDN (Anatrace). The Fab-γδ TCR complex was eluted using the wash buffer supplemented with 4 mM desthiobiotin. The co-precipitated proteins were analyzed by clear native PAGE and visualized using a fluorescence imager (Tanon MINI Space 3000).

### Native MS Analysis

The protein samples were buffer-exchanged into 1 M ammonium acetate (pH 7.0) using a Zeba 7K MWCO spin column (Thermo Fisher). The desalted protein samples were loaded into a home-made borosilicate capillary and electro-sprayed into an Orbitrap Ascend Tribrid mass spectrometer in positive mode (Thermo Fisher). The instrument was operated in intact-protein high-pressure mode. The typical MS settings were: 1,000 V electrospray voltage, 250 °C transfer tube temperature, Orbitrap resolution of 15,000, 150% RF lens, and 150-250 V source fragmentation. The mass spectra were processed using UniDec software (43).

### Mass Photometry

The molecular weight of proteins was determined using a Refeyn’s TwoMP mass photometry system (Refeyn Ltd.) following previously established protocols (44, 45). The samples were diluted in PBS (pH 7.4) or HBS (25 mM HEPES, pH 8.0, 150 mM NaCl), and the final concentration during measurement was 10-20 nM.

### N-Terminal Edman Sequencing

The UCHT1 IgG1 was separated into a light chain and a heavy chain on an SDS-PAGE gel under reducing conditions. The proteins were transferred to a methanol-activated PVDF membrane. After Ponceau S staining, the light chain (∼25 kDa) was excised. Then the membrane was washed with deionized water. The N-terminal sequence of the first six amino acids of the light chain was determined by automated Edman degradation using a Shimadzu PPSQ-33A protein sequencer (Shimadzu). The PTH-amino acid standard mixture was purchased from Wako (Japan).

### *De Novo* Antibody Sequencing by LC-MS/MS

The UCHT1 IgG1 was separated by SDS-PAGE. The corresponding gel bands were excised, reduced with dithiothreitol, and alkylated with iodoacetamide in the dark. The protein samples were digested with trypsin, chymotrypsin, Asp-N, and Lys-C at 37 °C overnight, respectively. The digested peptides were desalted using C18 StageTips, dried and reconstituted with 1% formic acid for LC-MS/MS analysis. The peptides were analyzed using a Vanquish Neo UHPLC coupled to an Orbitrap Exploris 480 mass spectrometer (Thermo Fisher Scientific). The peptides were separated with a binary buffer system on an in-house prepared capillary column (75 µm × 150 mm) packed with 1.9-µm, 120-Å ReproSil-Pur C18-AQ silica particles. Buffer A was 0.1% formic acid in 100% H_2_O and Buffer B was 0.1% formic acid in 80% acetonitrile with 20% H_2_O. Data were acquired in a data-dependent acquisition mode.

The raw data were first processed using PEAKS Studio 12.5 (Bioinformatics Solutions Inc). In the first stage, *de novo* sequencing was performed with precursor and fragment error tolerance of 10 ppm and 0.02 Da, respectively. Carbamidomethylation (Cys) was set as a fixed modification, and oxidation (Met) and deamidation (Asn/Gln) as variable modifications. The identified peptide sequences were exported along with their feature areas and positional confidence scores for subsequent full-length antibody reconstruction.

Fab-region sequences of the antibody were reconstructed from *de novo* LC-MS/MS data using the Stitch tool (https://github.com/snijderlab/stitch) (46). Peptide-spectrum matches generated by PEAKS software were aligned against a non-redundant set of human immunoglobulin V/J gene segment templates derived from the International ImMunoGeneTics information system (IMGT). Quality-and intensity-weighted consensus sequences were generated based on the alignment results. Analyses were performed using the default parameters (PEAKS Automatic Cleavage Confidence ≥ 50). All reconstruction results, including consensus sequences, sequence logos, and alignment coverage, were manually checked.

### Fab and Mutant Fab Production

The designs of the expression constructs and protein purification of UCHT1 and OKT3 Fabs were carried out essentially as described previously (9, 26, 27). The UCHT1 Fab sequence was established by integrating the previously reported UCHT1 sequence (9), *de novo* sequencing data of UCHT1 IgG1, N-terminal Edman sequencing of the UCHT1 IgG1 light chain, and structural information obtained from the cryo-EM structure of the UCHT1 Fab. The OKT3 Fab features a light chain composed of the Vκ domain of OKT3 fused to the murine Cκ domain, and a heavy chain comprising the V_H_ domain of OKT3 linked to the Cγ2a domain of murine IgG2a (26, 27). Both Fab chains were expressed under a modified cPTPRσ signal peptide (MGILPSPGMPALLSLVSLLSVLLMGCVA), followed by the chain sequences. A FLAG-tag followed by a 10×His-tag was engineered at the C-terminus of the Fab heavy chain. Constructs were cloned into the pcDNA3.1 expression vector.

When the cell density reached 3×10^6^ cells/ml, 0.5 mg each of the Fab heavy chain and Fab light chain plasmids and 5 mg of PEI (25 kDa, Polysciences) were mixed in 50 mL of fresh medium and incubated for 30 min before transfecting 1 L of Expi293F cells. Transfected cells were incubated at 37 °C and 5% CO_2_ for 24 h, then 10 mM sodium butyrate was added and incubated for an additional 72 h. Supernatant was collected after centrifugation (2000 rpm, 4 °C, 10 min), and incubated with Ni-NTA agarose resin (Lablead) for 12 h, which had been pre-equilibrated with 10 column volumes of lysis buffer (25 mM HEPES pH 8, 150 mM NaCl, 20 mM imidazole). After the mixture had been loaded onto the column, it was washed with 50 column volumes of washing buffer (25 mM HEPES pH 8, 150 mM NaCl, 50 mM imidazole). Bound Fab proteins were eluted with elution buffer containing 25 mM HEPES (pH 8), 150 mM NaCl, and 200 mM imidazole. Collected eluates were further purified using a Superdex 200 Increase 10/300 column (Cytiva) equilibrated with size-exclusion chromatography buffer (25 mM HEPES pH 8, 150 mM NaCl). In cellular functional assays, the Fab was buffer-exchanged into PBS using a 10 kDa MWCO filter (Amicon). The purity of the Fab was assessed by SDS-PAGE under reducing and non-reducing conditions.

### Lentivirus Production

To generate a TCRα-knockout Jurkat cell line, the *TRAC* locus was disrupted using CRISPR-Cas9 as previously described (47). The gRNA target sequences were: TRAC-gRNA-Top, 5’-TCCAGTTGGTGGCATTGCCG-3’; TRAC-gRNA-Bottom, 5’-CGGCAATGCCACCAACTGGA-3’ (48). For the Vγ9Vδ2 TCR-expressing Jurkat cell line, full-length Vγ9 and Vδ2 TCR chains linked by T2A peptide were subcloned into the pLVX-IRES-HygR vector. Lentiviral particles were produced by co-transfecting HEK293T cells with the transfer vector (either pLVX-IRES-HygR-Vγ9Vδ2 TCR or lentiCRISPRv2-Neo-gTRAC), along with the packaging plasmid psPAX2 and the envelope plasmid pMD2.G, using Lipofectamine 2000 (Thermo Fisher Scientific) according to the manufacturer’s instructions. At 48 h post-transfection, the virus was collected, filtered through a 0.45-μm syringe filter, and used for infection.

### Jurkat Cell Engineering

When Jurkat cells reached a density of 4×10^5^ cells/mL, the majority of the culture medium was replaced with unconcentrated lentiviral supernatant supplemented with 8 μg/mL polybrene. The cells were then centrifuged for 60 min at 2000 rpm. 24 h later, the lentiviral supernatant was largely replaced with fresh complete medium.

72 h post-infection, cells transduced with gTRAC lentivirus were selected in medium containing 1.6 mg/mL G418 (Biosharp). 7 days after selection, limiting dilution cloning was performed by resuspending cells at a density of 5 cells/mL and dispensing 100 μL per well into 96-well plates. After 3 weeks, individual clones were screened for TCRα protein expression by Western blot. Clone 1, which exhibited the lowest level of TCRα expression among all tested clones, was selected and designated as the TCRα-knockout (KO) Jurkat cell line.

To generate Vγ9Vδ2 TCR-expressing Jurkat cells (γδ TCR-Jurkat), the TCRα-KO Jurkat cells were transduced with Vγ9Vδ2 TCR viral supernatant and subjected to selection with 400 μg/mL hygromycin B (Biosharp) for 20 days. Surface expression of the γδ TCR-CD3 complex was assessed by flow cytometry.

### Cell Binding Dose Curves

5 × 10^5^ γδ TCR-Jurkat cells were incubated with serially diluted rFab for 30 min at 4 °C. The cells were then washed and incubated with Multi-rAb® CoraLite® Plus 647-conjugated goat anti-mouse recombinant secondary antibody (H+L) (Proteintech Group, Inc.,1:25 dilution) for 30 min at 4 °C, followed by an additional wash prior to flow cytometric analysis. Cell populations were first gated based on FSC-A versus SSC-A to exclude debris and subsequently on FSC-A versus FSC-H to exclude cell aggregates. K_D_ values were calculated using GraphPad Prism software.

### BRET Assays

BRET experiments were performed as described previously (8, 49). Plasmids were generated by fusing NanoLuc or mNeonGreen to the C-terminus of the Vδ2-T2A-Vγ9 TCR and cloning into pcDNA3.1. All BRET measurements were performed in phenol red-free cell culture medium. HEK293T cells (1.7 × 10^6^) seeded in 6-well plates were transfected with a mixture containing 1.5 µg CD3 plasmid, 1 µg Vδ2-T2A-Vγ9 TCR-mNeonGreen plasmid, 0.5 µg Vδ2-T2A-Vγ9 TCR-NanoLuc plasmid, and 15 µg PEI (25 kDa, Polysciences). 48 h post transfection, cells were plated in 96-well white flat bottom plates (2×10^4^ cells/well). After 24 h, cells were treated with PBS (vehicle control), 5 µg/mL IgG1, 3.33 µg/mL rFab, or 3.33 µg/mL mutant rFab (to obtain equimolar amounts). After incubation at room temperature for 30 min, furimazine was added to each sample to a final concentration of 3 μM. BRET signals were subsequently measured using a Varioskan Lux multimode microplate reader (Thermo Fisher Scientific, Monza, Italy), and the BRET ratio was calculated as Emission_510_/Emission_460_.

### T Cell Activation Assay

To assess CD69 upregulation, 2×10^5^ γδ TCR-Jurkat cells seeded in 96-well plates were treated with one of the following: PBS (vehicle control), 5 µg/mL IgG1, 3.33 µg/mL rFab, or 3.33 µg/mL mutant rFab (to obtain equimolar amounts). After incubation at 37 °C for 3 h, cells were harvested, washed with FPBS (PBS containing 1% FBS), and stained on ice for 40 min with PE-conjugated anti-human CD69 antibody (clone FN50, BioLegend) at a 1:25 dilution. Cells were then washed again with FPBS. Data were acquired on a CytoFLEX S (Beckman) and analyzed using FlowJo software (FlowJo LLC).

### Statistical Analysis

Statistical analyses were performed using GraphPad Prism 8. Data are presented as mean ± SD. Comparisons between two groups were analyzed by two-tailed unpaired Student’s *t*-test. For multiple-group comparisons, one-way ANOVA followed by Tukey’s multiple comparisons test was used. For untargeted lipidomic analyses of phospholipids, two-way ANOVA followed by Sidak’s multiple comparisons test was performed. For the vocanol plot of quantified phospholipids, two-sided unpaired Student’s *t*-tests was used. A *P* value < 0.05 was considered statistically significant. Significance is indicated as follows: ns, not significant; \**P* < 0.05; \*\**P* < 0.01; \*\*\**P* < 0.001; \*\*\*\**P* < 0.0001. *n* indicates the number of independent experiments.

## Data Availability

Cryo-EM maps and coordinates have been deposited in the EMDB and wwPDB, respectively, with accession numbers: EMD-61882 and PDB 9JXZ (γδ TCR); EMD-61883 and PDB 9JY0 (Fab-CD3γε-TCR); EMD-61884 and PDB 9JY1 (δε/δε Fab-TCR tetramer); EMD-61885 and PDB 9JY2 (Fab-CD3δε-TCR); EMD-61886 and PDB 9JY3 (δε/γε Fab-TCR tetramer); EMD-61887 and PDB 9JY4 (γε/γε Fab-TCR tetramer).

## Author Contributions

L.W. conceived the project. Z.L., Y.F., L.C., and Y.C. performed plasmid construction, protein purification, cell experiments, and lipidomic analysis. Z.L. prepared the cryo-EM samples. J.L. performed data collection. D.W., Y. Zhao, and Y. Zhang performed mass spectrometry. L.W. solved the cryo-EM structures. L.W., J.L., and Z.W. performed model building. L.W. and J.L. analyzed the structures. L.W., Z.L., and J.L. wrote the manuscript with input from Y. Zheng, J.W., D.W., and H.W.

## Acknowledgments

We are grateful to Danyang Li and Xiangning Li at the Core Facility of Wuhan University for Cryo-EM data collection services. We acknowledge Xiaolu Zhao and Tingfu Zhang at the College of Life Sciences of Wuhan University for lipidomic analysis. We thank Liang Cheng at Medical Research Institute of Wuhan University for gifting antibodies and providing guidance on flow cytometry experiments. We thank Songya Lyu from College of Life Sciences, Wuhan University, for the flow cytometry support. We also thank the Core Facility of the TaiKang Center for Life and Medical Sciences, Wuhan University for MS analysis. This work was supported by National Key R&D Program of China (2022YFA0912200), the Fundamental Research Funds for the Central Universities (2042025kf0012), Large-scale Instrument and Equipment Sharing Foundation of Wuhan University. D. W. is supported by Excellence Research Group Program of NSFC (210973074).

## Conflicts of Interest

The authors declare no conflicts of interest.

## Notes

### Competing Interest Statement

The authors have declared no competing interest.

### Summary of Updates

Title and abstract updated; funding information updated; reference to "Figs. 8 and 9" corrected to "Figs. S8 and S9"; "Author Contributions", "Acknowledgments", and "Conflicts of Interest" sections added; The title of Supplemental files updated.

