## Supplementary material for "Fab Dimerization Crosslinks and Activates the γδ T-Cell Receptor": Figures S1-S10, Tables S1-S3

### **Supporting Information for**
### Fab Dimerization Crosslinks and Activates the γδ T-Cell Receptor

Zezhi Li, Yihua Fan, Jing Li, Yuanyuan Zhao, Ligang Chen, Yiqing Chen, Zhiming Wang, Yumei Zheng, Yuxuan Zhang, Jun Wang, Hao Wu, Di Wu, and Longfei Wang

Corresponding author: Zezhi Li, Di Wu, and Longfei Wang

**This PDF file includes:**

Figures S1 to S10
Tables S1 to S3

#### Figures

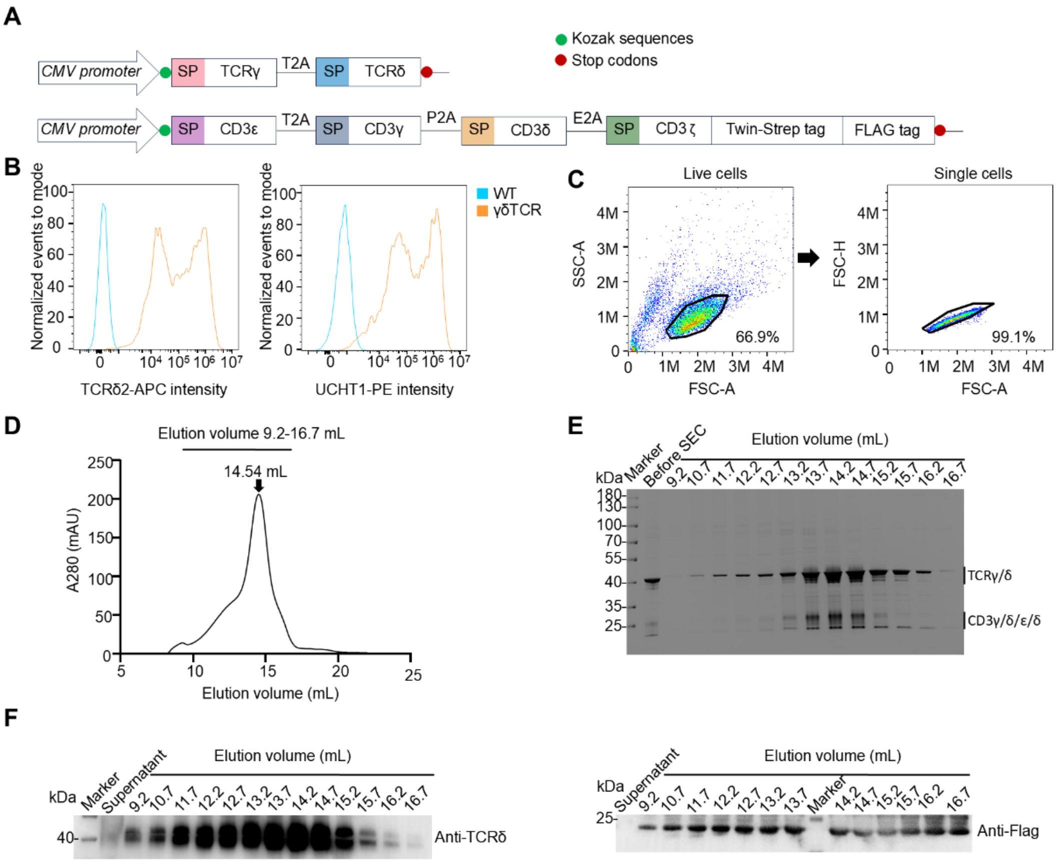


**Fig. S1. Purification of the fully assembled γδ TCR.** (***A***) Schematic showing the designs of the expression constructs. (***B***) Flow cytometric analyses of untransfected Expi293F cells (WT, blue) and Expi293F cells transfected with the TCR and CD3 plasmids (γδ TCR, orange). (***C***) Representative gating strategy for flow cytometry analysis. (***D***) Analytical size-exclusion chromatography of purified γδ TCR. (***E***) Reducing SDS-PAGE analysis of size-exclusion chromatography fractions. (***F***) Western blot analysis of size-exclusion chromatography fractions under reducing conditions.

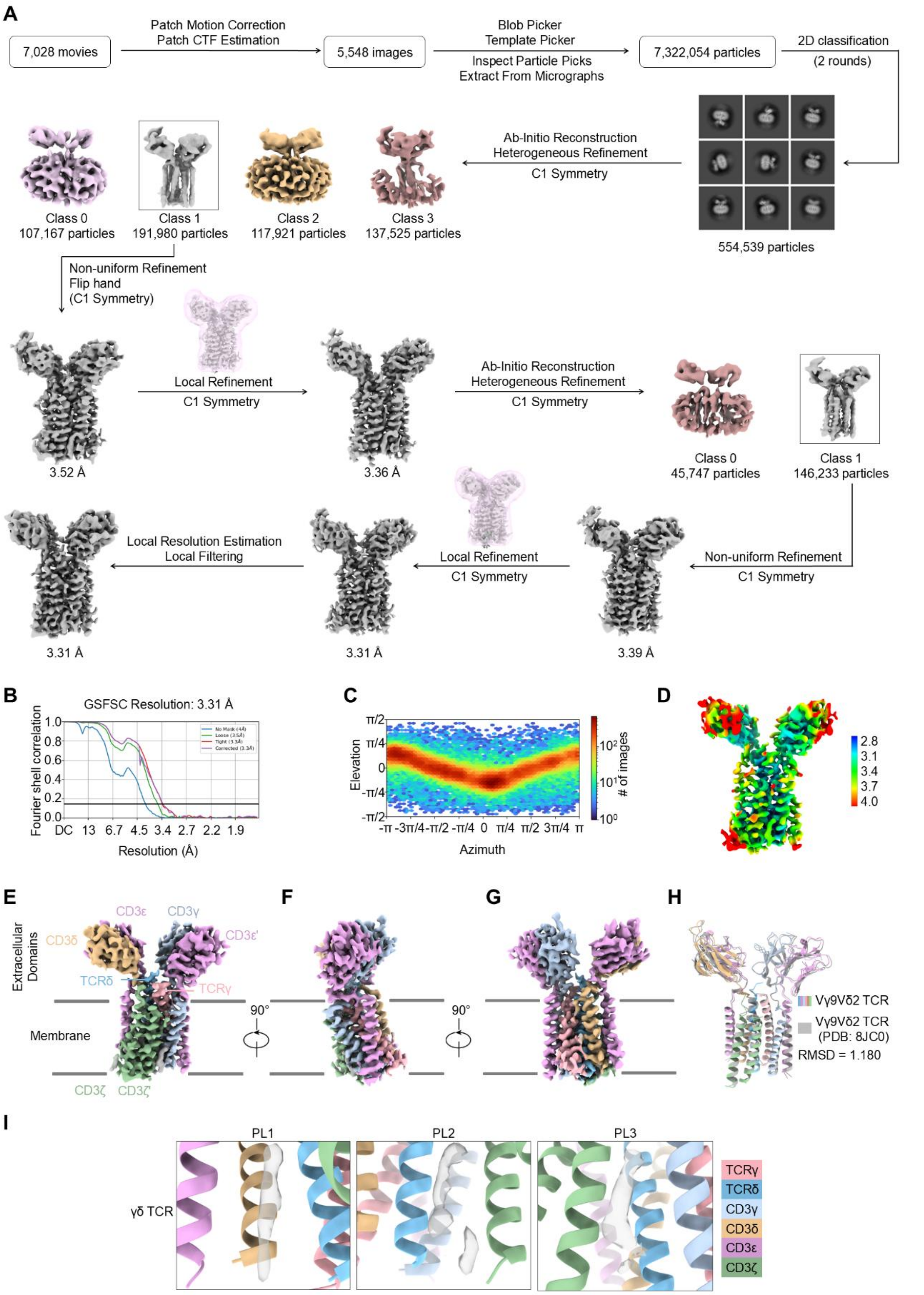


**Fig. S2. Cryo-EM analysis of the γδ TCR.** (***A***) Data processing workflow for the reconstruction of the γδ TCR. (***B***) Gold-standard FSC curves indicate an overall resolution of 3.31 Å at FSC=0.143.

(***C***) Orientation distribution plot of the 3D reconstruction of the γδ TCR. (***D***) Local resolution-filtered display of resolution (Å) colored from highest resolution (blue) to lowest resolution (red). (***E***) The Cryo-EM density of γδ TCR shown in side view. Subunits are colored as follows: TCRγ (pink), TCRδ (light sky blue), CD3γ (light steel blue), CD3δ (burly wood), CD3ε (plum), and CD3ζ (dark sea green). Membrane boundaries are indicated by black lines. (***F*** and ***G***) Cryo-EM densities of γδ TCR in two orthogonal views. (***H***) Overlaid ribbon diagram of Vγ9Vδ2 TCR (colored) and Vγ9Vδ2 TCR (PDB: 8JC0, gray). (***I***) Enlarged views of phospholipid (PL) binding interfaces in the γδ TCR.

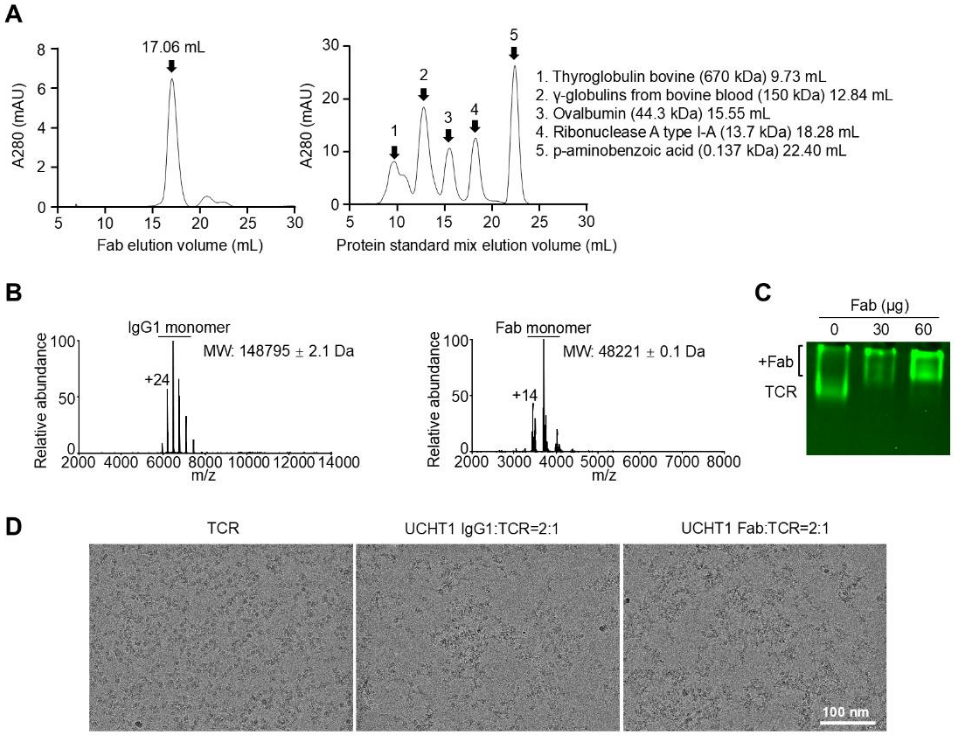


**Fig. S3. Cross-linking of γδ TCR complex by UCHT1 Fab.** (***A***) Analytical size-exclusion chromatography of Fab and protein standard mix. (***B***) Native mass spectrometry of UCHT1 IgG1 and Fab. (***C***) Binding of Fab to γδ TCR on Expi293F cells assessed by clear native PAGE. 6 × $10^7$ Expi293F cells expressing Twin-Strep-mNeonGreen-tagged γδ TCR were treated with 0, 30, or 60 µg Fab, lysed, and subjected to Twin-Strep tag-based enrichment. Eluates were analyzed by clear native PAGE and visualized using a fluorescence imager. (***D***) Representative original cryo-EM micrographs of the γδ TCR, UCHT1 IgG1-γδ TCR complex, and Fab-γδ TCR complex, which were imaged on a FEI Titan Krios G4 microscope operated at 300 kV.

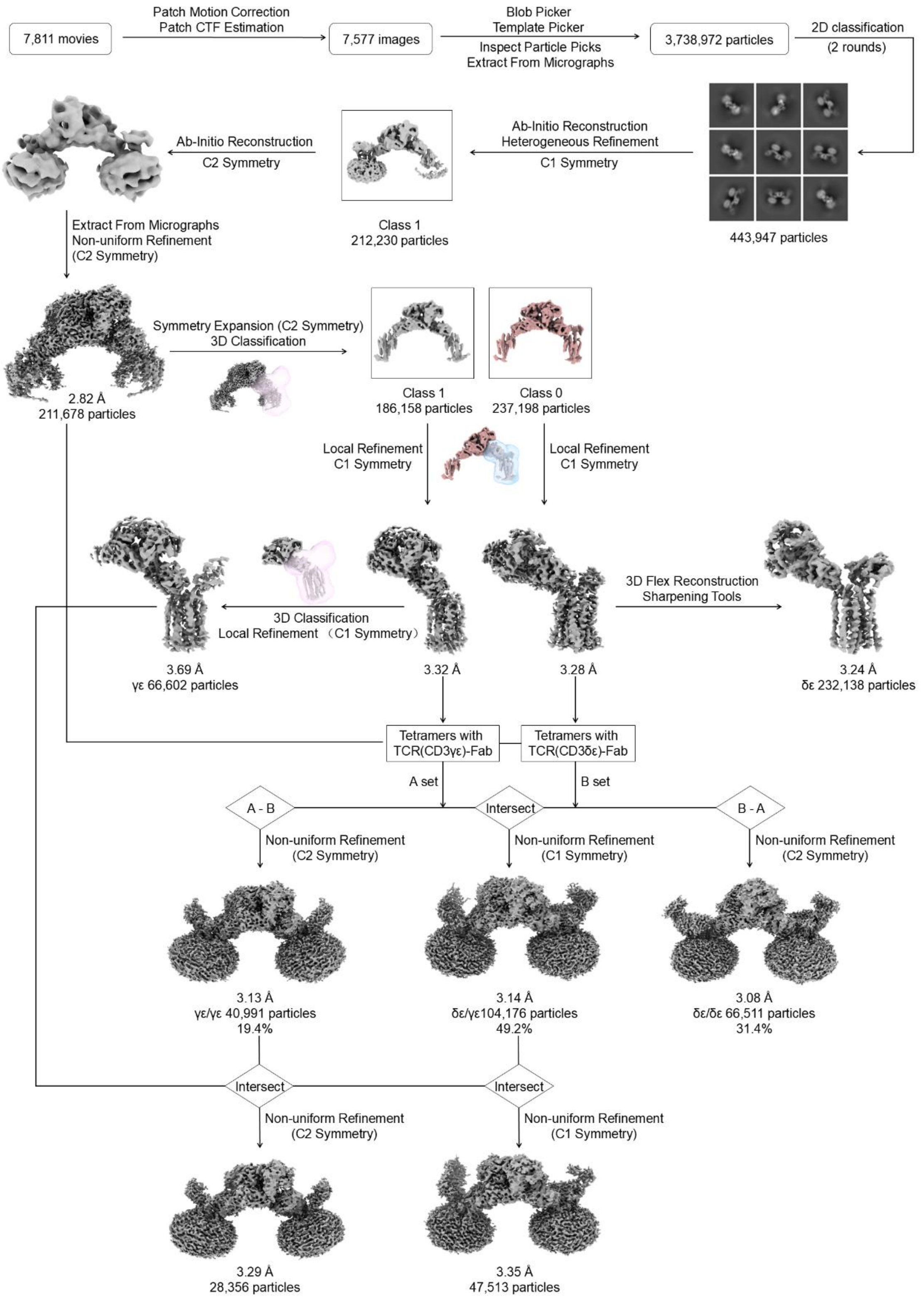


**Fig. S4. Cryo-EM data processing workflow for the reconstruction of UCHT1 Fab-γδ TCR complexes.**

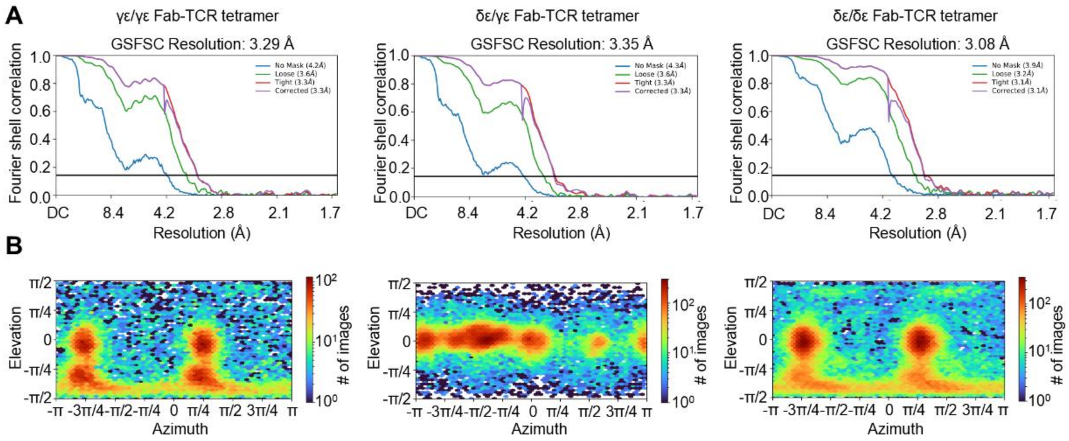


**Fig. S5. Cryo-EM analysis of the UCHT1 Fab-γδ TCR complexes.** (***A***) Gold standard FSC curves of γε/γε Fab-TCR tetramer (left), δε/γε Fab-TCR tetramer (middle), and δε/δε Fab-TCR tetramer (right), indicate overall resolution of 3.29 Å, 3.35 Å, and 3.08 Å at FSC=0.143, respectively. (***B***) Orientation distribution plot of the 3D reconstruction of the γε/γε Fab-TCR tetramer (left), δε/γε Fab-TCR tetramer (middle), and δε/δε Fab-TCR tetramer (right).

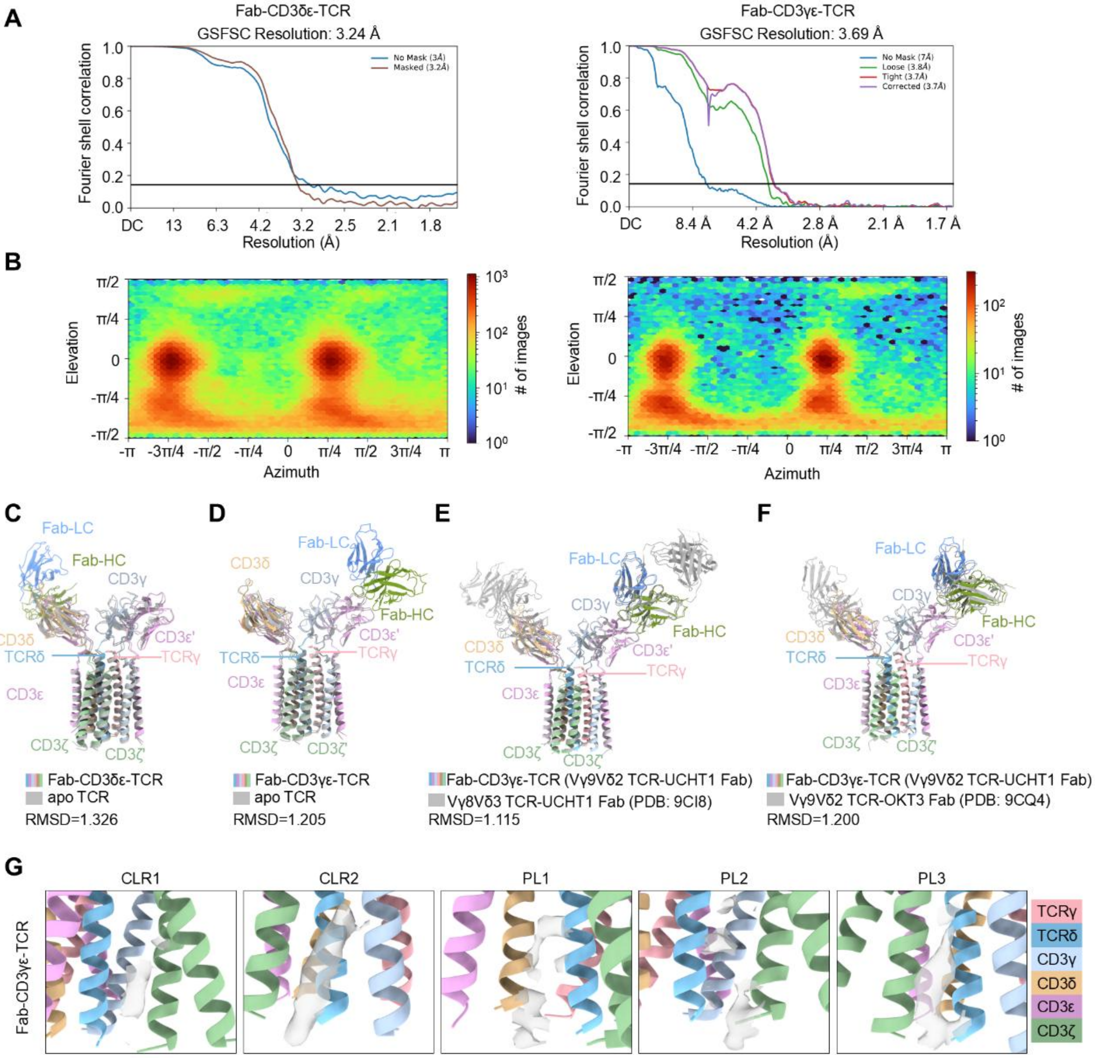


**Fig. S6. Cryo-EM analysis and conformation changes of Fab-TCR complexes.** (***A***) Gold standard FSC curves of Fab-CD3δε-TCR (left) and Fab-CD3γε-TCR (right), indicate overall resolution of 3.24 Å and 3.69 Å at FSC=0.143, respectively. (***B***) Orientation distribution plot of the 3D reconstruction of the Fab-CD3δε-TCR (left) and Fab-CD3γε-TCR (right). (***C***) Overlaid ribbon diagram of the Fab-CD3δε-TCR (colored) and apo TCR (gray). (***D***) Overlaid ribbon diagram of the Fab-CD3γε-TCR (colored) and apo TCR (gray). (**E**) Overlaid ribbon diagram of the Vγ9Vδ2 TCR-UCHT1 Fab (colored) and Vγ8Vδ3 TCR-UCHT1 Fab (PDB: 9CI8, gray). (**F**) Overlaid ribbon diagram of the Vγ9Vδ2 TCR-UCHT1 Fab (colored) and Vγ9Vδ2 TCR-OKT3 Fab (PDB: 9CQ4, gray). (**G**) Enlarged views of cholesterol (CLR) and phospholipid (PL) binding interfaces in the Fab-CD3γε-TCR. TCRγ (pink), TCRδ (light sky blue), CD3γ (light steel blue), CD3δ (burly wood), CD3ε (plum), and CD3ζ (dark sea green).

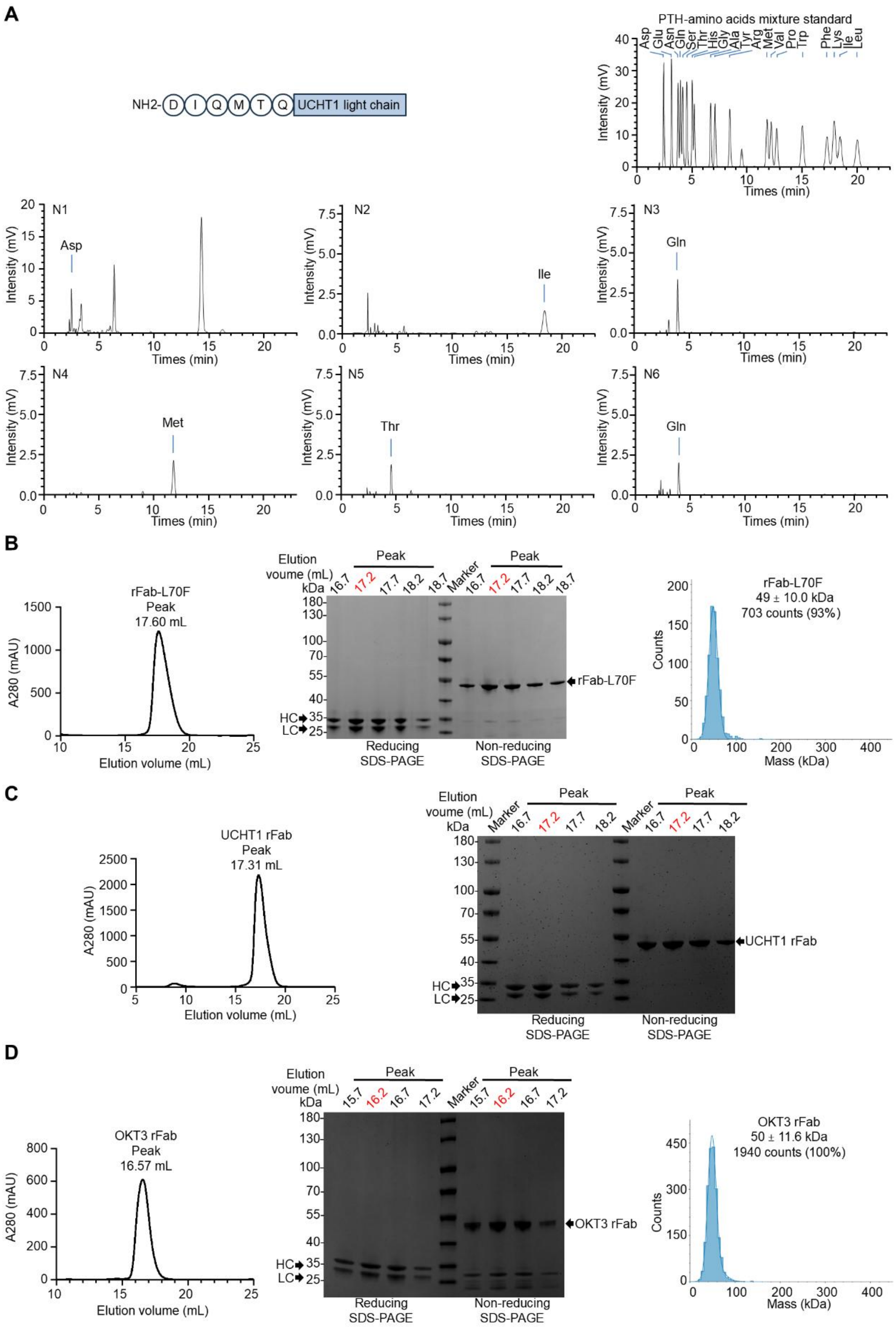


**Fig. S7. N-terminal Edman sequencing of the UCHT1 IgG1 light chain and purification of**

**wild-type and mutant rFabs.** (***A***) Edman sequencing of N-terminus of the UCHT1 IgG1 light chain. (***B***) Analytical size-exclusion chromatography (left), SDS-PAGE analysis of size-exclusion chromatography fractions (middle), and mass photometry (right) of rFab-L70F (L70F in the heavy chain). (***C***) Analytical size-exclusion chromatography (left) and SDS-PAGE analysis of size-exclusion chromatography fractions (right) of UCHT1 rFab. (***D***) Analytical size-exclusion chromatography (left), SDS-PAGE analysis of size-exclusion chromatography fractions (middle), and mass photometry (right) of OKT3 rFab. In ***B***-***D***, size-exclusion chromatography fractions used for subsequent experiments are highlighted in red in the SDS-PAGE panels.

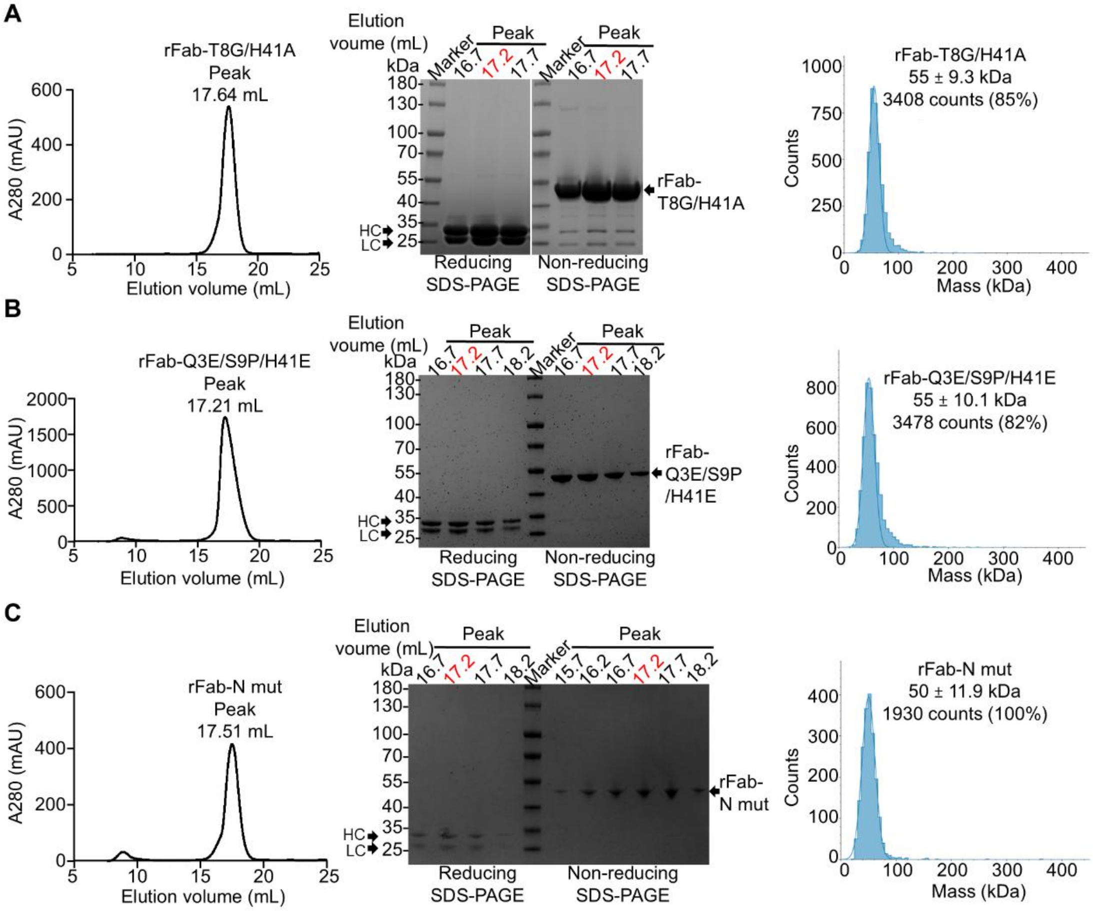


**Fig. S8. Purification of dimer-disrupting rFab mutants.** (***A***) Analytical size-exclusion chromatography (left), SDS-PAGE analysis of size-exclusion chromatography fractions (middle), and mass photometry (right) of rFab-T8G/H41A (T8G in the light chain and H41A in the heavy chain). (***B***) Analytical size-exclusion chromatography (left), SDS-PAGE analysis of size-exclusion chromatography fractions (middle), and mass photometry (right) of rFab-Q3E/S9P/H41E (Q3E/S9P in the light chain and H41E in the heavy chain). (***C***) Analytical size-exclusion chromatography (left), SDS-PAGE analysis of size-exclusion chromatography fractions (middle), and mass photometry (right) of rFab-N mut (deleting residues $D_1IQ_3$ and replacing residues $T_7TSS_{10}$ with GSGS in the light chain). In ***A***-***C***, size-exclusion chromatography fractions used for subsequent experiments are highlighted in red in the SDS-PAGE panels.

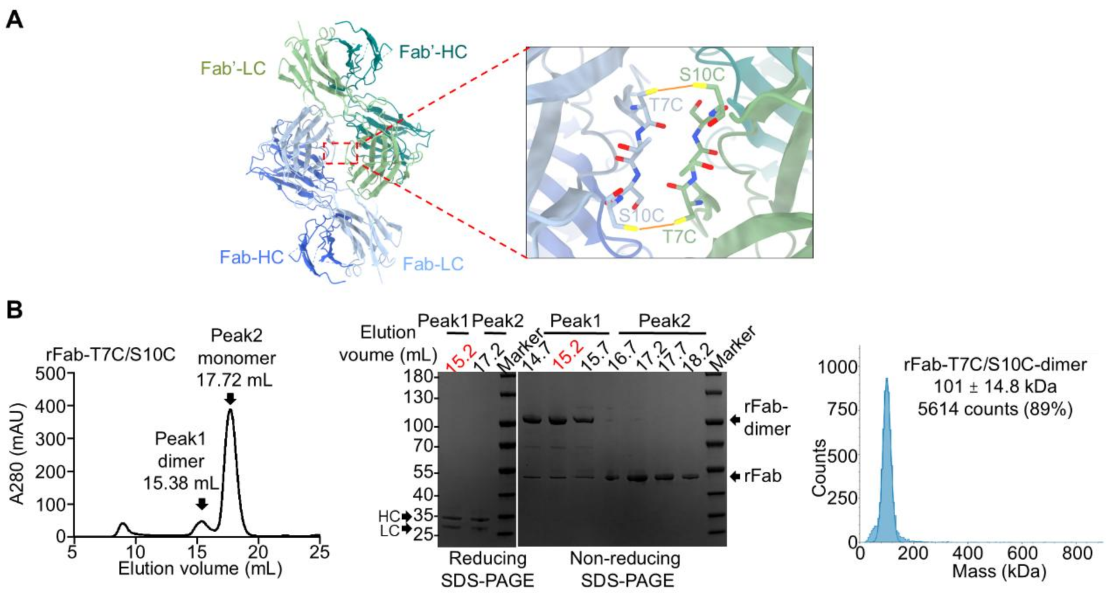


**Fig. S9. Purification of dimer-stabilizing rFab mutants.** (***A***) Schematic representation of the rFab-T7C/S10C-dimer (T7C/S10C in the light chain) ribbon diagram. One Fab heavy and light chains are colored in royal blue and light steel blue respectively, and another one Fab heavy and light chains are colored in teal and dark sea green respectively. Orange lines denote disulfide bonds. (***B***) Analytical size-exclusion chromatography (left), SDS-PAGE analysis of size-exclusion chromatography fractions (middle), and mass photometry (right) of rFab-T7C/S10C-dimer. Size-exclusion chromatography fractions used for subsequent experiments are highlighted in red in the SDS-PAGE panels.

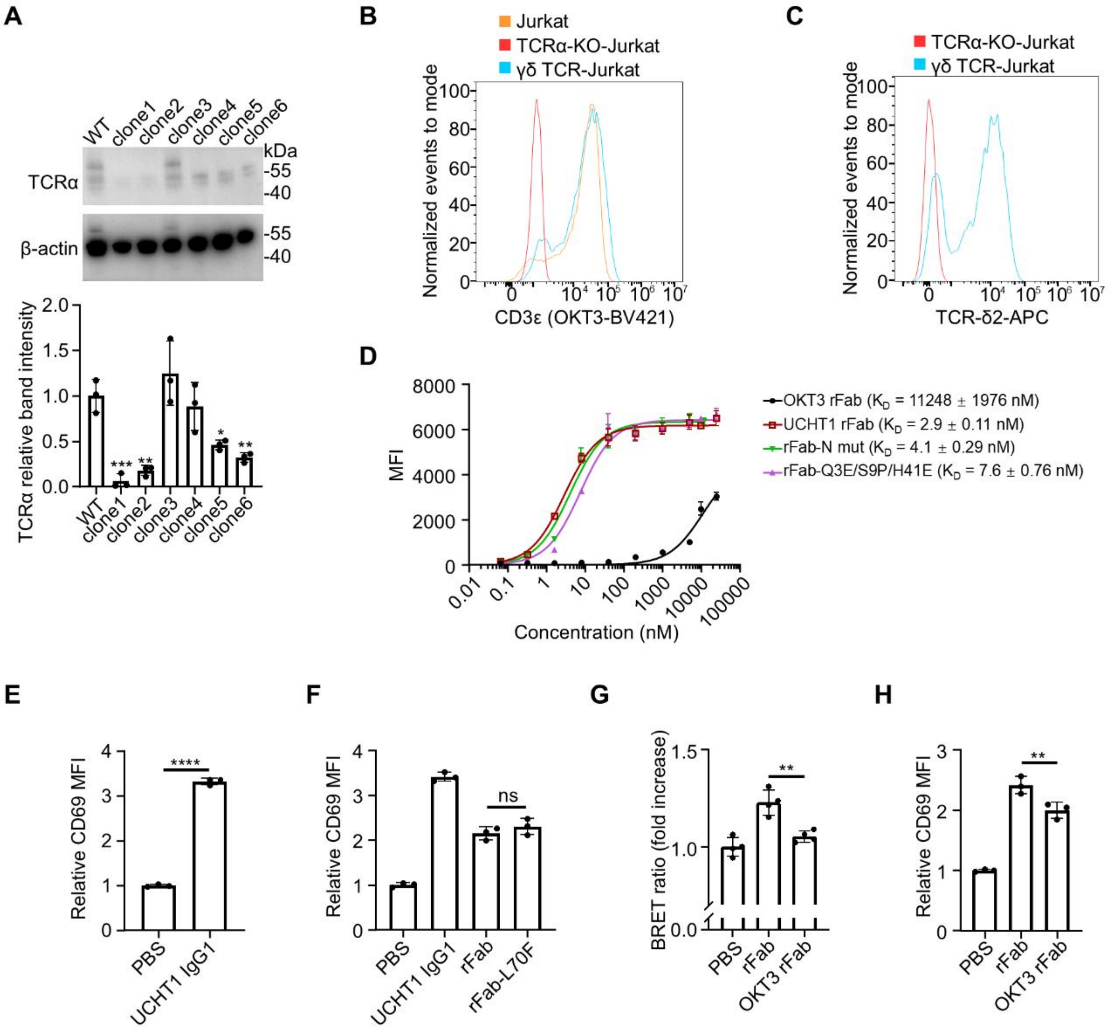


**Fig. S10. Fab dimerization enhances TCR activation.** (***A***) Screening of TCRα-KO Jurkat monoclonal cell lines by Western blot for TCRα ablation (*n* = 3 biologically independent experiments; mean ± SD). (***B***) Flow cytometric analysis of CD3 expression in Jurkat, TCRα-KO-Jurkat, and γδ TCR-Jurkat cells. (***C***) Flow cytometric analysis of TCRδ expression in Jurkat, TCRα-KO-Jurkat, and γδ TCR-Jurkat cells. (***D***) Binding of rFabs and rFab mutants to γδ TCR-Jurkat cells assessed by flow cytometry (*n* = 3 biologically independent experiments; mean ± SD). MFI indicates median fluorescence intensity. (***E*** and ***F***) γδ TCR-expressing Jurkat cells were stimulated with PBS (vehicle control), UCHT1 IgG1, rFab, or rFab-L70F for 3 h, followed by staining with PE-conjugated anti-human CD69 antibody for flow cytometric analysis of CD69 upregulation (*n* = 3 biologically independent experiments; mean ± SD). MFI, median fluorescence intensity. (***G***) BRET ratios in Expi293F cells co-expressing γδ TCR-NanoLuc and γδ TCR-mNeonGreen upon treatment with PBS, UCHT1 rFab, or OKT3 rFab (*n* = 4 biologically independent experiments; mean ± SD). (***H***) γδ TCR-expressing Jurkat cells were stimulated with PBS, UCHT1 rFab, or OKT3 rFab for 3 h, followed by staining with PE-conjugated anti-human CD69 antibody for flow cytometric analysis of CD69 upregulation (*n* = 3 biologically independent experiments; mean ± SD). *P* values were calculated using one-way ANOVA followed by Tukey's multiple comparisons test (***A*** and ***F***-***H***) or by two-tailed unpaired *t*-test (***E***). Statistical significance: ns, not significant; **P* < 0.05; ***P* < 0.01; ****P* < 0.001; *****P* < 0.0001.

**Tables**

**Table S1. Sequences of the protein constructs used in this study.**

| Name | Protein sequence |
|---|---|
| TCRγ9 | MLSLLHTSTLAVLGALCVYGAGHLEQPQISSTKTLSKTARLECVVSGIT<br>ISATSVYWYRERPGEVIQFLVSISYDGTVRKESGIPSGKFEVDRIPETS<br>TSTLTIHNVEKQDIATYYCALWEAQQELGKKIKVFGPGTKLIITDKQLDA<br>DVSPKPTIFLPSIAETKLQKAGTYLCLLEKFFPDVIKIHWEEKKSNTILG<br>SQEGNTMKTNDTYMKFSWLTVPEKSLDKEHRCIVRHENNKNGVDQEI<br>IFPPIKTDVITMDPKDNCSKDANDTLLLQLTNTSAYYMYLLLLLKSVVYF<br>AIITCCLLRRTAFCCNGEKS |
| TCRδ2 | MQRISSLIHLSLFWAGVMSAIELVPEHQTVPVSIGVPATLRCSMKGEAI<br>GNYYINWYRKTQGNTMTFIYREKDIYGPGFKDNFQGDIDIAKNLAVLKI<br>LAPSERDEGSYYCACDTLGMGGEYTDKLIFGKGTRVTVEPRSQPHTK<br>PSVFVMKNGTNVACLVKEFYPKDIRINLVSSKKITEFDPAIVISPSGKYN<br>AVKLGKYEDSNSVTCSVQHDNKTVHSTDFEVKTDSTDHVKPKETENT<br>KQPSKSCHKPKAIVHTEKVNMMSLTVLGLRMLFAKTVAVNFLLTAKLFF<br>L |
| CD3ε | MQSGTHWRVLGLCLLSVGVWGQDGNEEMGGITQTPYKVSISGTTVIL<br>TCPQYPGSEILWQHNDKNIGGDEDDKNIGSDEDHLSLKEFSELEQSG<br>YYVCYPRGSKPEDANFYLYLRARVCENCMEMDVMSVATIVIVDICITG<br>GLLLLVYYWSKNRKAKAKPVTRGAGAGGRQRGQNKERPPPVPNPDY<br>EPIRKGQRDLYSGLNQRRI |
| CD3γ | MEQGKGLAVLILAIILLQGTLAQSIKGNHLVKVYDYQEDGSVLLTCDAE<br>AKNITWFKDGKMIGFLTEDKKKWNLGSNAKDPRGMYQCKGSQNKSK<br>PLQVYYRMCQNCIELNAATISGFLFAEIVSIFVLAVGVYFIAGQDGVRQ<br>SRASDKQTLLPNDQLYQPLKDREDDQYSHLQGNQLRRN |
| CD3δ | MEHSTFLSGLVLATLLSQVSPFKIPIEELEDRVFVNCNTSITWVEGTVG<br>TLLSDITRLDLGKRILDPRGIYRCNGTDIYKDKESTVQVHYRMCQSCV<br>ELDPATVAGIIVTDVIATLLLALGVFCFAGHETGRLSGAADTQALLRND<br>QVYQPLRDRDDAQYSHLGGNWARNK |
| CD3ζ | MKWKALFTAAILQAQLPITEAQSFGLLDPKLCYLLDGILFIYGVILTALFL<br>RVKFSRSADAPAYQQGQNQLYNELNLGRREEYDVLDKRRGRDPEMG<br>GKPQRRKNPQEGLYNELQKDKMAEAYSEIGMKGERRRGKGHDGLY<br>QGLSTATKDTYDALHMQALPPRSA |
| UCHT1 rFab-LC | DIQMTQTTSSLSASLGDRVTISCRASQDIRNYLNWYQQKPDGTVKLLI<br>YYTSRLHSGVPSKFSGSGSGTDYSLTISNLEQEDIATYFCQQGNTLP<br>WTFAGGTKLEIKRADAAPTVSIFPPSSEQLTSGGASVVCFLNNFYPKDI<br>NVKWKIDGSERQNGVLNSWTDQDSKDSTYSMSSTLTLTKDEYERHN<br>SYTCEATHKTSTSPIVKSFNRNEC |
| UCHT1 rFab-HC | EVQLQQSGPELVKPGASMKISCKASGYSFTGYTMNWVKQSHGKNLE<br>WMGLINPYKGVSTYNQKFKDKATLTVDKSSSTAYMELLSLTSEDSAVY<br>YCARSGYYGDSDWYFDVWGAGTTVTVSSAKTTPPSVYPLAPGSAAQ<br>TNSMVTLGCLVKGYFPEPVTVTWNSGSLSSGVHTFPAVLQSDLYTLS |

| | |
|---|---|
| | SSVTVPSSPRPSETVTCNVAHPASSTKVDKKIVPRDCGGGSDYKDDDDKGGGGSHHHHHHHHHH |
| UCHT1 rFab-H-L70F | EVQLQQSGPELVKPGASMKISCKASGYSFTGYTMNWVKQSHGKNLEWMGLINPYKGVSTYNQKFKDKATFTVDKSSSTAYMELLSLTSEDSAVYYCARSGYYGDSDWYFDVWGAGTTVTVSSAKTTPPSVYPLAPGSAAQTNSMVTLGCLVKGYFPEPVTVTWNSGSLSSGVHTFPAVLQSDLYTLSSSVTVPSSPRPSETVTCNVAHPASSTKVDKKIVPRDCGGGSDYKDDDDKGGGGSHHHHHHHHHH |
| UCHT1 rFab-L-mut | MTQGSGSLSASLGDRVTISCRASQDIRNYLNWYQQKPDGTVKLLIYYTSRLHSGVPSKFSGSGSGTDYSLTISNLEQEDIATYFCQQGNTLPWTFAGGTKLEIKRADAAPTVSIFPPSSEQLTSGGASVVCFLNNFYPKDINVKWKIDGSERQNGVLNSWTDQDSKDSTYSMSSTLTLTKDEYERHNSYTCEATHKTSTSPIVKSFNRNEC |
| UCHT1 rFab-L-Q3E/S9P | DIEMTQTTPSLSASLGDRVTISCRASQDIRNYLNWYQQKPDGTVKLLIYYTSRLHSGVPSKFSGSGSGTDYSLTISNLEQEDIATYFCQQGNTLPWTFAGGTKLEIKRADAAPTVSIFPPSSEQLTSGGASVVCFLNNFYPKDINVKWKIDGSERQNGVLNSWTDQDSKDSTYSMSSTLTLTKDEYERHNSYTCEATHKTSTSPIVKSFNRNEC |
| UCHT1 rFab-H-H41E | EVQLQQSGPELVKPGASMKISCKASGYSFTGYTMNWVKQSEGKNLEWMGLINPYKGVSTYNQKFKDKATLTVDKSSSTAYMELLSLTSEDSAVYYCARSGYYGDSDWYFDVWGAGTTVTVSSAKTTPPSVYPLAPGSAAQTNSMVTLGCLVKGYFPEPVTVTWNSGSLSSGVHTFPAVLQSDLYTLSSSVTVPSSPRPSETVTCNVAHPASSTKVDKKIVPRDCGGGSDYKDDDDKGGGGSHHHHHHHHHH |
| UCHT1 rFab-L-T8G | DIQMTQTGSSLSASLGDRVTISCRASQDIRNYLNWYQQKPDGTVKLLIYYTSRLHSGVPSKFSGSGSGTDYSLTISNLEQEDIATYFCQQGNTLPWTFAGGTKLEIKRADAAPTVSIFPPSSEQLTSGGASVVCFLNNFYPKDINVKWKIDGSERQNGVLNSWTDQDSKDSTYSMSSTLTLTKDEYERHNSYTCEATHKTSTSPIVKSFNRNEC |
| UCHT1 rFab-H-H41A | EVQLQQSGPELVKPGASMKISCKASGYSFTGYTMNWVKQSAGKNLEWMGLINPYKGVSTYNQKFKDKATLTVDKSSSTAYMELLSLTSEDSAVYYCARSGYYGDSDWYFDVWGAGTTVTVSSAKTTPPSVYPLAPGSAAQTNSMVTLGCLVKGYFPEPVTVTWNSGSLSSGVHTFPAVLQSDLYTLSSSVTVPSSPRPSETVTCNVAHPASSTKVDKKIVPRDCGGGSDYKDDDDKGGGGSHHHHHHHHHH |
| UCHT1 rFab-L-T7C/S10C | DIQMTQCTSCLSASLGDRVTISCRASQDIRNYLNWYQQKPDGTVKLLIYYTSRLHSGVPSKFSGSGSGTDYSLTISNLEQEDIATYFCQQGNTLPWTFAGGTKLEIKRADAAPTVSIFPPSSEQLTSGGASVVCFLNNFYPKDINVKWKIDGSERQNGVLNSWTDQDSKDSTYSMSSTLTLTKDEYERHNSYTCEATHKTSTSPIVKSFNRNEC |
| OKT3 rFab-L | QIVLTQSPAIMSASPGEKVTMTCSASSSVSYMNWYQQKSGTSPKRWIYDTSKLASGVPAHFRGSGSGTSYSLTISGMEAEDAATYYCQQWSSNPFTFGSGTKLEIKRADAAPTVSIFPPSSEQLTSGGASVVCFLNNFYPKDINVKWKIDGSERQNGVLNSWTDQDSKDSTYSMSSTLTLTKDEYERHNSYTCEATHKTSTSPIVKSFNRNEC |

| | |
|---|---|
| OKT3 rFab-H | QVQLQQSGAELARPGASVKMSCKASGYTFTRYTMHWVKQRPGQGL<br>EWIGYINPSRGYTNYNQKFKDKATLTTDKSSSTAYMQLSSLTSEDSAV<br>YYCARYYDDHYCLDYWGQGTTLTVSSAKTTAPSVYPLAPVCGGTTG<br>SSVTLGCLVKGYFPEPVTLTWNSGSLSSGVHTFPAVLQSDLYTLSSSV<br>TVTSSTWPSQSITCNVAHPASSTKVDKKIEPRGGGSDYKDDDDKGGG<br>GSHHHHHHHHHH* |

**Table S2. Sequences of the UCHT1 Fab by *de novo* sequencing of UCHT1 IgG1.**

| Name | Protein sequence |
|---|---|
| UCHT1 LC | DIQMTQTTSSLSASLGDRVTISCRASQDIRNYLNWYQQKPDGTVKLLIYYT<br>SRLHSGVPSKFSGSGSGTDYSLTISNLEQEDIATYFCQQGNTLPWTFAGGT<br>KLEIKRADAAPTVSIFPPSSEQLTSGGASVVCFLNNFYPKDINVKWKIDGSE<br>RQNGVLNSWTDQDSKDSTYSMSSTLTLTKDEYERHNSYTCEATHKTSTSP<br>IVKSFNRNEC |
| UCHT1 HC | EVQLQQSGPELVKPGASMKISCKASGYSFTGYTMNWVKQSHGKNLEWM<br>GLINPYKGVSTYNQKFKDKATLTVDKSSSTAYMELLSLTSEDSAVYYCARS<br>GYYGDSDWYFDVWGAGTTVTVSSAKTTPPSVYPLAPGSAAQTNSMVTLG<br>CLVKGYFPEPVTVTWNSGSLSSGVHTFPAVLQSDLYTLSSSVTVPSSPRPS<br>ETVTCNVAHPASSTKVDKKIVPRDC |

#### Table S3. Cryo-EM data collection, refinement and validation statistics.

| | γδ TCR | γε/γε Fab-TCR tetramer | δε/γε Fab-TCR tetramer | δε/δε Fab-TCR tetramer | Fab-CD3δε-TCR | Fab-CD3γε-TCR |
|---|---|---|---|---|---|---|
| EMDB ID | 61882 | 61887 | 61886 | 61884 | 61885 | 61883 |
| PDB ID | 9JXZ | 9JY4 | 9JY3 | 9JY1 | 9JY2 | 9JY0 |
| **Data collection and processing** | | | | | | |
| Magnification | 105,000 | 105,000 | | | | |
| Voltage (kV) | 300 | 300 | | | | |
| Electron exposure (e–/Å$^2$) | 50 | 50 | | | | |
| Defocus range (μm) | -1.5 to -1.8 | -1.5 to -1.8 | | | | |
| Pixel size (Å) | 0.84 | 0.824 | | | | |
| Symmetry imposed | C1 | C2 | C1 | C2 | C1 | C1 |
| Initial particle images (no.) | 7,322,054 | 3,738,972 | 3,738,972 | 3,738,972 | 3,738,972 | 3,738,972 |
| Final particle images (no.) | 146,233 | 28,356 | 47,513 | 66,511 | 232,138 | 66,602 |
| Map resolution (Å) | 3.31 | 3.29 | 3.35 | 3.08 | 3.24 | 3.69 |
| FSC threshold | 0.143 | 0.143 | 0.143 | 0.143 | 0.143 | 0.143 |
| Map resolution range (Å) | 2.8 to 7.1 | 2.8 to 6.9 | 1.8 to 7.0 | 1.8 to 5.9 | 1.8 to 5.8 | 3.0 to 7.4 |
| **Refinement** | | | | | | |
| Initial model used (PDB) | 6JXR | 9JXZ | 9JXZ | 9JXZ | 9JXZ | 9JXZ |
| Model resolution (Å) | 3.6 | 4.1 | 3.7 | 3.8 | 3.5 | 3.9 |
| FSC threshold | 0.5 | 0.5 | 0.5 | 0.5 | 0.5 | 0.5 |
| Model resolution range (Å) | 3.6 to 50 | 4.1 to 50 | 3.7 to 50 | 3.8 to 50 | 3.5 to 50 | 3.9 to 50 |
| Map sharpening *B* factor (Å$^2$) | -103.8 | -62.8 | -56.1 | -75.3 | -75.2 | -70.8 |
| Model composition | | | | | | |
| Non-hydrogen atoms | 4846 | 15642 | 15638 | 15642 | 6515 | 6521 |
| Protein residues | 597 | 2002 | 1998 | 1996 | 818 | 822 |
| Ligands | 5 | 0 | 2 | 4 | 3 | 1 |
| *B* factors (Å$^2$) | | | | | | |
| Protein | 71.43 | 93.2 | 90.28 | 77.45 | 108.5 | 101.81 |
| Ligand | 48.97 | 0 | 94.74 | 93.23 | 86.45 | 97.65 |
| R.m.s. deviations | | | | | | |
| Bond lengths (Å) | 0.003 | 0.003 | 0.003 | 0.003 | 0.005 | 0.004 |
| Bond angles (°) | 0.582 | 0.749 | 0.746 | 0.775 | 0.782 | 0.897 |
| Validation | | | | | | |
| MolProbity score | 2.41 | 2.25 | 2.43 | 2.53 | 2.52 | 2.57 |
| Clash score | 10.35 | 12.97 | 14.32 | 15.86 | 17.54 | 17.83 |
| Poor rotamers (%) | 3.58 | 2.39 | 4.11 | 6.18 | 5.26 | 3.17 |
| Ramachandran plot | | | | | | |
| Favored (%) | 92.94 | 95.17 | 95.67 | 96.49 | 96.48 | 93.27 |
| Allowed (%) | 7.06 | 4.73 | 4.18 | 3.51 | 3.52 | 6.73 |
| Disallowed (%) | 0 | 0.1 | 0.15 | 0 | 0 | 0 |
